# The Evolutionary History of Wild, Domesticated, and Feral *Brassica oleracea* (Brassicaceae)

**DOI:** 10.1101/2021.04.06.438638

**Authors:** Makenzie E. Mabry, Sarah D. Turner-Hissong, Evan Y. Gallagher, Alex C. McAlvay, Hong An, Patrick P. Edger, Jonathan D. Moore, David A. C. Pink, Graham R. Teakle, Chris J. Stevens, Guy Barker, Joanne Labate, Dorian Q. Fuller, Robin G. Allaby, Timothy Beissinger, Jared E. Decker, Michael A. Gore, J. Chris Pires

## Abstract

Understanding the evolutionary history of crops, including identifying wild relatives, helps to provide insight for designing new approaches in crop breeding efforts. Cultivated *Brassica oleracea* has intrigued researchers for centuries due to its wide diversity in forms, which include cabbage, broccoli, cauliflower, kale, kohlrabi, and Brussels sprouts. Yet, the evolutionary history of this species remains understudied. With such different vegetables produced from a single species, *B. oleracea* is a model organism for understanding the power of artificial selection. Persistent challenges in the study of *B. oleracea* include conflicting hypotheses regarding domestication and the identity of the closest living wild relative. Using a diversity panel of 224 accessions, which represents 14 different *B. oleracea* crop types and nine potential wild progenitor species, we integrate phylogenetic and population genetic techniques with ecological niche modeling, archaeological, and literary evidence to examine relationships among cultivars and wild relatives to clarify the origin of this horticulturally important species. Our analyses point to the Aegean endemic *B. cretica* as the closest living relative of cultivated *B. oleracea*, supporting an origin of cultivation in the Eastern Mediterranean region. Additionally, we identify several feral lineages, suggesting that cultivated plants of this species are able to revert to a wild-like state with relative ease. By expanding our understanding of the evolutionary history in *B. oleracea*, these results contribute to a growing body of knowledge on crop domestication that will facilitate continued breeding efforts including adaptation to changing environmental conditions.

## INTRODUCTION

> “*Greek legend has it that the cabbage sprung from where Zeus’ sweat hit the ground*.”
>
> — -*N. D. Mitchell (1976)*

A key tenet of evolutionary and plant biology is understanding how plants respond and adapt to changes in environmental conditions, which requires leveraging genotypic diversity and understanding the connections between genotype and phenotype. Crop wild relatives (CWRs) provide pools of allelic diversity that at one time were shared through a common ancestor with cultivated relatives. Although Vavilov recognized the potential of CWRs in the early 1900s (Vavilov 1926), advances in genomics and genome editing techniques have enabled scientists to better realize the potential of CWRs as a source of diversity and novel traits for the improvement of cultivated populations (Prohens et al. 2017; Li et al. 2018; Fernie and Yan 2019; Khoury et al. 2020; Turner-Hissong et al. 2020). Yet these scientific advancements are hindered in that we still have not identified the CWRs of many important crop species. One such species, *Brassica oleracea* L., has an unclear evolutionary history due to taxonomic confusion and the lack of genetic and archaeological evidence.

The horticultural crop *B. oleracea* has played an important role in global food systems for centuries, providing a source of leaf and root vegetables, fodder, and forage (Shyam et al. 2012). When first introduced to the species, Darwin drew many parallels between his theory of natural selection and the cultivation practices that led to the varied forms of this plant (Darwin and Gray 1868). While many people may recognize that various dog breeds are descended from wolves, most people are surprised to learn that the domesticated forms of *B. oleracea*, broccoli (var. *italica*), Brussels sprouts (var. *gemmifera*), cabbage (var. *capitata*), cauliflower (var. *botrytis*), kale (var. *acephala*), and kohlrabi (var. *gongylodes*) are the same species. Although just six major crop types comprise the majority of the U.S. market (Agricultural Marketing Service, Market News Reports; www.ams.usda.gov), the global market for *B. oleracea* crops was around 70.1 million metric tons, in terms of production for 2019 (The Food and Agriculture Organization; www.fao.org). Outside these six major cultivars there exists at least 12 additional cultivated crop types (***SI Appendix*, Table S1**). These include lesser known varieties such as Chinese white kale or Cantonese gai-lan (Mandarian Mandarin *Jiè lán* 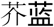; **var.** *alboglabra*), a leafy vegetable with florets, romanesco (var. *botrytis*) with unique fractal patterned curds, and walking stick kale (var. *longata*), which grows 6-12 feet in height.

Compared to other crops, surprisingly little is known about the progenitor species and origin of domesticated *B. oleracea*. Primary challenges in identifying the progenitor species include the number of wild species that share a single cytodeme and are interfertile with *B. oleracea* (2n = 18 chromosomes; similar genomic organization; referred to as the “C genome”), the corresponding confusion surrounding taxonomic relationships, and conflicting evidence regarding the center of origin. Wild relatives that share the C genome with domesticated *B. oleracea* include *Brassica bourgeaui, Brassica cretica, Brassica hilarionis, Brassica incana, Brassica insularis, Brassica macrocarpa, Brassica montana, Brassica rupestris*, and *Brassica villosa*. Throughout the literature, many of these species have been referred to by alternative names or have multiple subspecies. For example, *B. cretica* is described as having either three subspecies (subsp. *aegea, cretica*, and *laconica*) (Snogerup et al. 1990) or only two (subsp. *cretica* and *nivea (Gustafsson et al. 1976*)). The taxonomic confusion is perhaps best highlighted by L. H. Bailey, who stated that “*Some of these plants appear to be more confused in literature than in nature*” (Bailey 1930)). The progenitor species of *B. oleracea* is further obscured by the presence of weedy, cabbage-like plants along the coastline of western Europe (England, France, and Spain), which are sometimes referred to as *B. oleracea* var. *sylvestris*. The role of these weedy populations in the domestication of *B. oleracea* is unclear, with some studies suggesting these coastal wild populations represent the progenitor species (Snogerup et al. 1990; Song et al. 1990), and others identifying these wild forms as plants that escaped cultivation (Mitchell 1976; Mitchell and Richards 1979).

Given the uncertainty surrounding wild relatives and weedy populations, researchers have proposed numerous hypotheses for the progenitor species of *B. oleracea* (**Table 1**). Hypotheses range from a single domestication with a single progenitor species (Song et al. 1990; Allender et al. 2007) to multiple domestications arising from multiple progenitor species (de Candolle 1855; Neutrofal 1927; Lizgunova 1959; Helm 1963; Snogerup 1980; Heaney et al. 1987; Song et al. 1988; Swarup and Brahmi 2005). Findings that point to a single origin of domestication have proposed different wild species as the progenitor (Snogerup et al. 1990; Song et al. 1990; Hodgkin 1995; Maggioni et al. 2018). For instance, (Neutrofal 1927) suggested that *B. montana* was the progenitor of cabbages and that *B. rupestris* was the progenitor of kohlrabi, while (Schulz 1936) identified *B. cretica* as the progenitor of only cauliflower and broccoli. (Helm 1963) proposed a triple origin in which a single progenitor species gave rise to cauliflower, broccoli, and sprouting broccoli, while kale and Brussels sprouts were derived from another unknown wild species, and that all other crop forms were derived from a third unknown wild species. (Snogerup 1980) proposed that cabbages were derived from wild *B. oleracea*, kales were derived from both *B. rupestris* and *B. incana*, and that Chinese white kale was derived specifically from *B. cretica* subsp. *nivea*.

**Table 1.** Wild species which have been proposed as progenitor species for *B. oleracea* crop types. Specific location in parentheses if indicated by the author.

| **Cultivar** | **Wild relative** | **Author** |
| --- | --- | --- |
| Broccoli | *B. oleracea* | Linnaeus |
|  | *B. oleracea* | Hedrick (1919) * |
|  | *B. oleracea* | Giles (1941) ** |
|  | *B. montana* | Hegi (1919) |
|  | *B. oleracea* (from Italy) | Giles (1941) |
|  | *B. cretica* | Gates (1953) |
|  | *B. oleracea* and *B. alboglabra* | Song et al. (1990) |
| Brussels sprouts | *B. oleracea* | Linnaeus |
|  | *B. oleracea* (western Europe) | Gates (1953) |
|  | *B. oleracea* (western Europe) | Snogerup (1980) |
|  | *B. oleracea* and *B. alboglabra* | Song et al. (1990) |
| Cabbage | *B. oleracea* | Linnaeus |
|  | *B. oleracea* | de Candolle (1824) |
|  | *B. oleracea* | Hedrick (1919) * |
|  | *B. oleracea* | Bailey (1930) |
|  | *B. montana* | Hegi (1919) |
|  | *B. oleracea* (western Europe) | Gates (1953) |
|  | *B. oleracea* (western Europe) | Snogerup (1980) |
|  | *B. oleracea* and *B. alboglabra* | Song et al. (1990) |
| Cauliflower | *B. oleracea* | Linnaeus |
|  | *B. oleracea* | de Candolle (1824) |
|  | *B. oleracea* | Bailey (1930) |
|  | *B. montana* | Hegi (1919) |
|  | *B. cretica* | Schulz (1936) |
|  | *B. oleracea* (from Cyprus) | Giles (1941) |
|  | *B. cretica* | Gates (1953) |
|  | *B. oleracea* and *B. alboglabra* | Song et al. (1990) |
|  | *B. cretica* | Tutin et al. (1964) |
| Kale | *B. oleracea* | Linnaeus |
|  | *B. oleracea* | Hedrick (1919) * |
|  | *B. oleracea* | Bailey (1930) |
|  | *B. montana* | Hegi (1919) |
|  | *B. montana* | Netroufal (1927) |
|  | *B. oleracea* (western Europe) | Gates (1953) |
|  | *B. cretica*, *B. incana*, *B. rupestris* | Snogerup (1980) |
|  | *B. incana* and *B. insularia* | Hosaka et al. (1990) |
|  | *B. oleracea* and *B. alboglabra* | Song et al. (1990) |
| Kohlrabi | *B. oleracea* | Linnaeus |
|  | *B. rupestris* | Netroufal (1927) |
|  | Unknown Mediterranean species | Gates (1953) |
|  | *B. oleracea* and *B. alboglabra* | Song et al. (1990) |

\* edited observations by Sturtevant in the late 19th century, \*\* referring to Prof Buckman's experiment

Due to the lack of consensus on the progenitor species, the center of origin for *B. oleracea* has also remained obscure. One hypothesis is that domesticated *B. oleracea* originated in England from weedy *B. oleracea* populations, with early cultivated forms brought to the Mediterranean, where selection for many of the early crop types occurred (Hodgkin 1995). Other studies point specifically to Sicily, which boasts a large diversity of wild relatives, as the center of domestication (Schiemann 1932; Lizgunova 1959). This conforms with the observations of Nikolai Vavilov (Vavilov 1951) that plants tended to be domesticated in a finite number of global centers of diversity, which include the Mediterranean. Most recently, linguistic and literary evidence provided support for domestication in the Eastern Mediterranean, where there is a rich history of expressions related to the usage and cultivation of *B. oleracea* crop types in early Greek and Latin literature (Maggioni et al. 2018).

Using a diversity panel of 224 accessions that includes 14 cultivar types and nine wild relatives, representing the largest and most diverse collection of this species and its wild relatives to date, we integrate phylogenomics, population genomics, species distribution modeling, archaeological, and literary sources to clarify the taxonomy, identify the closest living wild relative, and provide insight on the origin of domestication for *B. oleracea*.

## RESULTS

### Phylogeny and population clustering distinguish wild and feral populations

Sampling of *B. oleracea* cultivars included eight types of kales, five types of cabbages, Brussels sprouts, broccoli, cauliflower, Romanesco (var. *botrytis*), and kohlrabi (***SI Appendix*, Table S2**). Together, these cultivated types accounted for 188 of the 224 total samples. The remaining 36 samples included previously identified wild relatives: putatively wild *B. oleracea, B. cretica, B. incana, B. montana, B. hilarionis, B. insularis, B. macrocarpa, B. rupestris*, and *B. villosa*. The phylogenetic reconstruction of all 224 samples using SNPhylo (Lee et al. 2014) recovered several well-supported clades with greater than 70% bootstrap support, although overall support was generally poor (less than 70% bootstrap support), especially along the backbone. Chinese white kale, broccoli, cauliflower, romanesco, kohlrabi, curly kale, Brussels sprouts, *B. rupestris, B. macrocarpa*, and *B. insularis* were all recovered as monophyletic. Cabbages were also monophyletic, but with only 55% bootstrap support. Seven cultivars (collards, tronchuda kale, savoy cabbage, perpetual kale, red cabbage, and marrow cabbage) were found throughout the tree as polyphyletic assemblages. Several wild samples were recovered within the cultivar clade, including two samples of *B. cretica* (196, 199), one sample *of B. montana* (222), and all samples of putatively wild *B. oleracea* (175, 176, 177; sample names in red; **Fig. 1**). We also recovered a monophyletic group in the cultivar clade consisting of five samples of three wild species, *B. incana* (205, 208, 209), *B. villosa* (233), and *B. cretica* (195), labeled ‘wild C - clade 2’ (for wild samples with the C genome). Many of these “wild” samples also share most or all of their ancestry with cultivars. At K = 2, in our fastSTRUCTURE analyses (Raj et al. 2014), where the marginal likelihood is maximized, samples clustered together as either cultivars or wild individuals (**Fig. 1**). We find that two samples of *B. incana* (204, 207), which are sister to all cultivated samples, share 100% of their ancestry with cultivated types, as do two samples of *B. cretica* (196, 199), one sample *of B. montana* (222), and all three samples of putatively wild *B. oleracea* (175, 176, 177). Together with the placement in the phylogeny, these analyses indicate that these are not truly wild samples, but represent feral types. Our newly identified wild C - clade 2 also shows mixed wild and cultivar ancestry, which was also observed for one sample of tronchuda kale (30). At K = 3, clusters of broccoli, cauliflower, and Chinese white kale separate from other cultivated types and at K = 4, Chinese white kale separates from broccoli and cauliflower.

**Figure 1:**
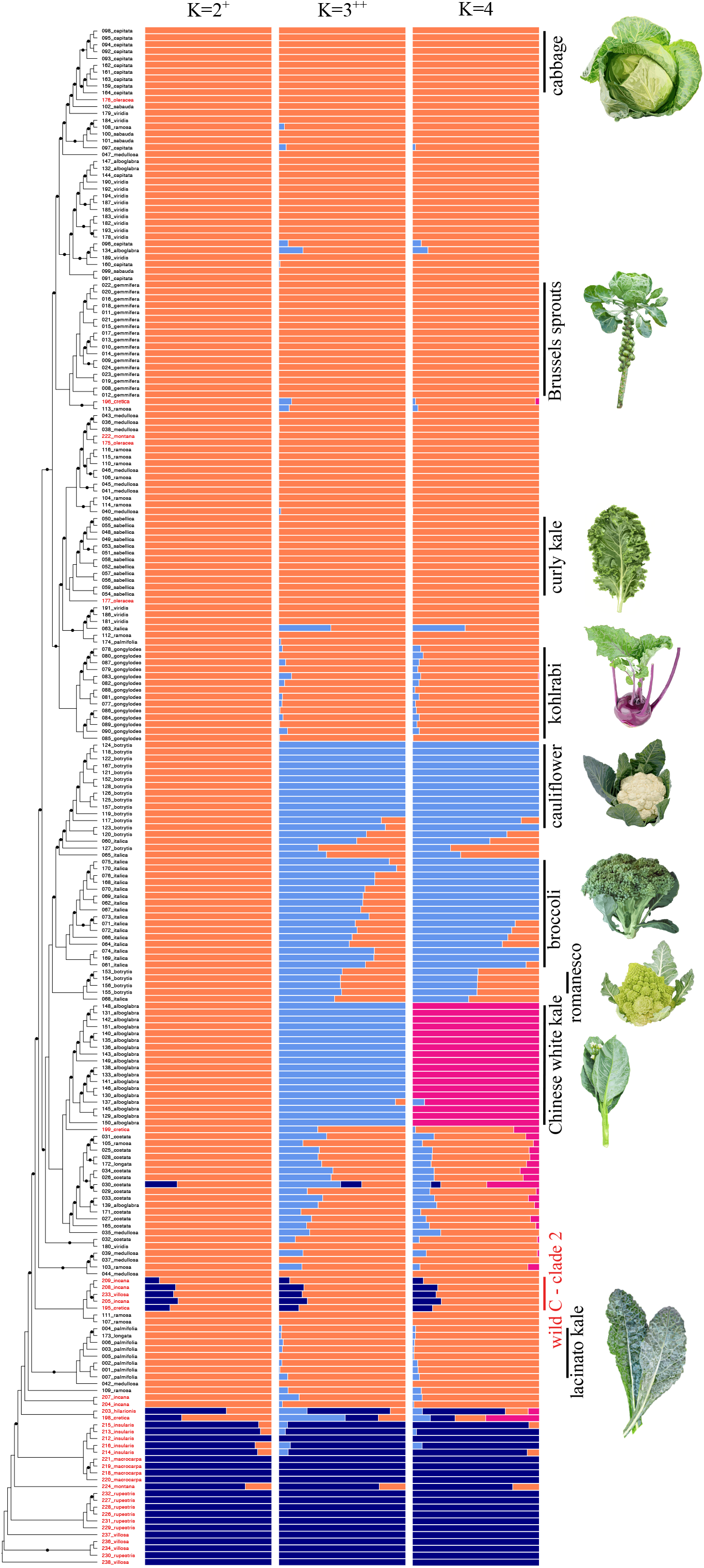
Demographics and population structure for 224 samples of cultivated Brassica oleracea (n=188) and wild C genome species (n = 36). (Left) Individual sample phylogeny with putatively wild samples labeled in red and black dots indicating bootstrap values less than 70%. (Middle) Ancestry proportions for K = 2 to K = 4 as inferred from fastSTRUCTURE; K=2 maximizes marginal likelihood (+) and K=3 best explains structure in the data (++). (Right) Major clades and illustrations of corresponding crop types. Illustrations by Andi Kur.

Principal component analysis (PCA) also separated cultivars from most wild samples (**Fig. 2A & B; *SI Appendix*, Fig. S1A**). The PC1 axis distinguishes wild species from cultivars and the PC2 axis separates wild C - clade 2 from all other wild species (triangles with black outlines). While one sample of *B. cretica* (198) clusters closest to cultivated types, samples of *B. incana*, not in wild C - clade 2, along with one sample of *B. montana* (222), two samples of *B. cretica* (196, 199), and all three samples of *B. oleracea* (175, 176, 177) cluster with the cultivars, corroborating the phylogenetic analyses. To further investigate the clustering patterns of *B. cretica* to cultivars, we included four additional wild-collected samples of two *B. cretica* subspecies (A and B = subsp. *nivea*, C and D = subsp. *cretica;* **Fig. 2A & B**; ***SI Appendix*, Fig. S1A;** labeled SRA in figure legend; (Kioukis et al. 2020)). Adding these samples supports the results of other studies that *B. cretica*, as a species, is very diverse. While sample C does not group with other *B. cretica* samples using the PC1 axis, the PC2, PC3, and PC4 axes show much tighter clustering among the four wild-collected samples and one of our samples of *B. cretica* (198), indicating that our *B. cretica* (198) sample is an informative representative of wild-collected *B. cretica* (**Fig. 2A & B; *SI Appendix*, Fig. S1A**).

**Figure 2:**
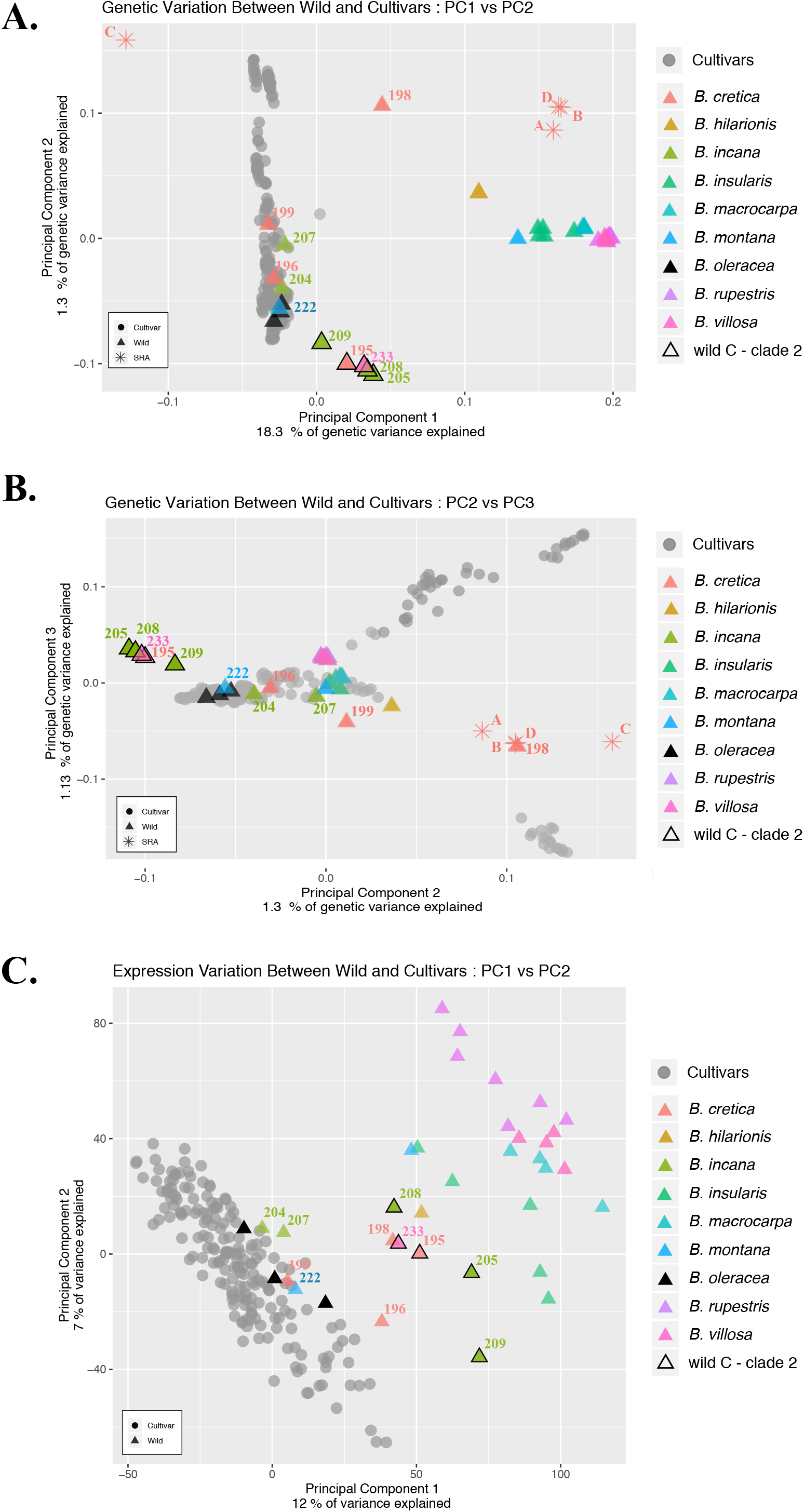
Principal Component Analysis (PCA) of SNPs and expression profiles. **A)** Genetic variation PCA of PC1 vs PC2, **B)** Genetic variation PC2 vs PC3, and **C)** Expression profile PCA for PC1 vs PC2 of wild and cultivar samples. Triangles = wild samples, circles = cultivars. Triangles with black outlines = wild C - clade 2 samples with species identification indicated by color. Wild-collected *B. cretica* samples from (Kioukis et al. 2020) indicated by asterisks, labeled as SRA.

For crop samples, estimates of inbreeding coefficients from PCAngsd (Meisner and Albrechtsen 2018) roughly matched expectations for the frequency of heterozygotes under Hardy-Weinberg equilibrium, while inbreeding coefficients for wild species suggest excess homozygosity (***SI Appendix*, Fig. S2**), possibly reflecting cultivation practices for germplasm management and the relative isolation of wild populations (i.e. small effective population size), respectively. Identified feral samples (*B. cretica* −196, 199; *B. incana* - 204, 207; *B. montana* - 222; and wild *B. oleracea* −175, 176, 177) show patterns of heterozygosity similar to crop samples, as does the four samples of *B. cretica* from (Kioukis et al. 2020). Our wild C - clade 2 is found with patterns of excess homozygosity more similar to other wild taxa.

### Domestication is also reflected in the transcriptome

Using expression profiles (transcript abundances) of 51,438 genes for our original 224 samples, we tested if cultivars and wild samples would still cluster separately based on the transcriptome. Overall, results and clustering patterns were similar to analyses using SNPs, with the axes of PC1 and PC2 separating most wild species from cultivars (**Fig. 2C; *SI Appendix*, Fig. S1B & C**). We again found the same samples *of B. incana* (204, 207), *B. cretica* (196, 199), *B. montana* (222), and *B. oleracea* (175, 176, 177) clustering with the cultivars, but in expression analyses wild C - clade 2 clustered with the other wild samples, rather than separately. Hierarchical clustering of the expression profiles recovered similar patterns with two major groups: wild and cultivated, again with wild C - clade 2 clustering with the other wild samples (***SI Appendix*, Fig. S3**). While most cultivar groups were not recovered as unique clusters, there were a few exceptions. Brussels sprouts, Chinese white kale, and curly kale all formed fairly distinct clades, which corresponds to what we know about their growth habit. Since RNA was collected at the 7th leaf-stage, before substantial morphological differentiation occurs between cultivars, it is not too surprising that they do not cluster distinctively by cultivar. However, curly kale is almost immediately visually distinguishable from other cultivars in that the first true leaves are already curly. Brussels sprouts are also identifiable at this early growing stage as they have orbicular leaves rather than the more lanceolate leaves of other cultivars. While Chinese white kale leaves look more similar to other cultivars, they are annuals and grow much more rapidly which may be responsible for its clustering separately.

To identify modules of genes that might be driving the observed clustering patterns, we used WGCNA (Langfelder and Horvath 2008). We found that 48 modules, ranging in size from 35,981 to 34 genes, provided the best fit for the data (***SI Appendix*, Table S3**). To assess what types of biological processes were overrepresented in these modules, we used syntenic *Arabidopsis thaliana* genes and performed a GO analysis through PANTHER v. 16.0 (Mi et al. 2021). Overlap *of B. oleracea* with *A. thaliana* genes ranged from 17% to 98.3%, perhaps indicating that either some modules are more conserved while others are unique to *B. oleracea*. Modules which were more conserved between the two species included genes related to herbivory defense compound production (secondary metabolite biosynthetic process, phenylpropanoid biosynthetic and metabolic processes), wound formation (suberin biosynthetic processes), and wax formation (wax biosynthetic and metabolic processes) likely correlated to the characteristic glaucous leaves of cultivated *B. oleracea* (***SI Appendix*, Table S4**).

### Species tree and admixture inference indicate *Brassica cretica* is the closest living wild relative

Given the results of population clustering using both SNPs and expression profiles, we further interrogated the species level relationships between wild relatives and cultivar groups by resolving the backbone of the phylogeny. Using the PoMo model (Schrempf et al. 2016) as implemented in IQ-Tree (Nguyen et al. 2015) and only including samples representing monophyletic groups as determined in the sample-level phylogeny, we found strong support for *B. cretica* as the closest living wild relative to cultivated *B. oleracea* (**Fig. 3A**). The current distribution of *B. cretica* occurs throughout the Eastern Mediterranean, primarily in Greece, highlighting a potential origin of domestication (**Fig. 3B**). Another suggested wild relative, *B. incana*, is strongly supported as belonging to the cultivar clade, sister to lacinato kale. This result supports our other findings that *B. incana* is not a wild assemblage, but rather feral. Within cultivars, several expected relationships were recovered: collards and cabbage as sister lineages (Song et al. 1988; Farnham 1996), with Brussels sprouts sister to both; cauliflower and broccoli as sister clades (Song et al. 1988; Stansell et al. 2018), with romanesco sister to both; and Chinese white kale as sister to all other cultivars, agreeing with recent literature (Cheng et al. 2016; Stansell et al. 2018).

**Figure 3:**
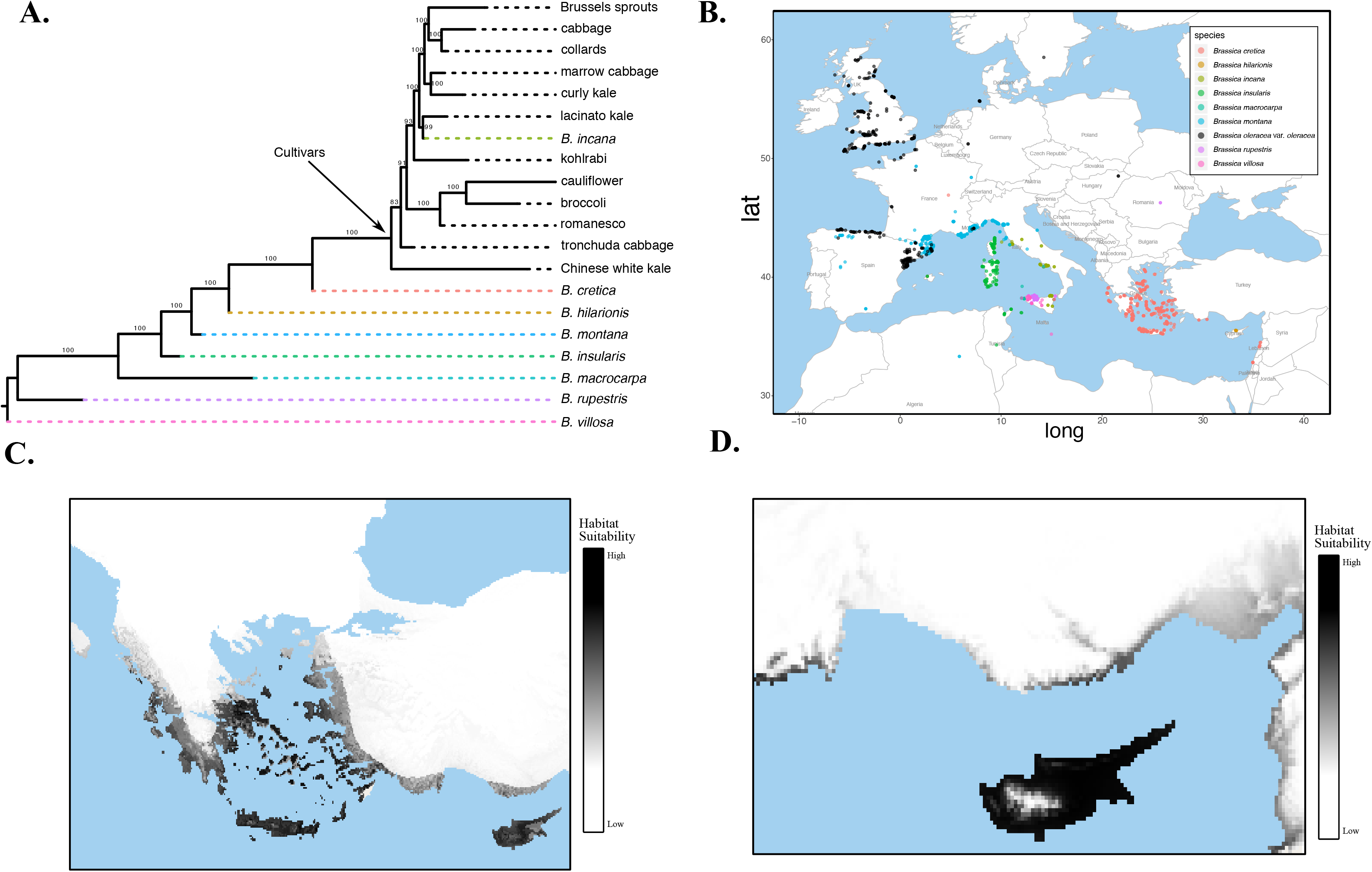
Species tree with current distribution and historical environmental niche modeling. **A)** Species tree of wild and cultivar samples. Bootstrap support indicated above branches. **B)** Current species distribution of wild relatives. **C)** Suitable habitat for *B. cretica* and **D)** *B. hilarionis* during the late-Holocene.

With the overall species relationships resolved, we aimed to tease apart the evolutionary history of the wild samples that clustered within the cultivar clade. Specifically, we asked if any of the identified feral samples were the products of admixture using TreeMix (Pickrell and Pritchard 2012). While the tree model without any migration edges explained 87.3% of the variance in the dataset, sequentially adding migration events to the tree resulted in five migrations events explaining 92% of the variation (**Fig. 4A; *SI Appendix*, Fig. S4**). Adding a single migration edge resulted in an admixture event from *B. cretica* (198) to a clade of [Chinese white kale + tronchuda cabbage]. To further test this event, we used four-population (*f*4) tests for treeness as implemented in TreeMix, where a significant non-zero value indicates the presence of gene flow (Reich et al. 2009; Pickrell and Pritchard 2012); **Fig. 4B**). While the tree [[tronchuda cabbage, kohlrabi],[*B. cretica* (198), *B. hilarionis]]* showed no significant evidence of gene flow (*f*4 = 0.0008, Z = 1.094), replacing tronchuda cabbage with Chinese white kale indicated significant gene flow from *B. cretica* (198) to Chinese white kale (*f*4 = −0.0055, Z = −5.113). This result was further verified when adding a second migration edge, as the migration edge only included Chinese white kale, but the direction was reversed (from Chinese white kale to *B. cretica* (198)). The second event, from kohlrabi to a presumably feral sample of *B. cretica* (199), was supported by *f4* tests, with the tree [[kohlrabi, *B. cretica* (196)],[*B. cretica* (199), marrow cabbage]] indicating significant evidence of gene flow from kohlrabi to *B. cretica* (199) (*f4* = 0.012, Z = 10.5). This migration event is also seen phenotypically, as *B. cretica* (199) has a swollen stem when grown to maturity. No significant evidence of gene flow was found when substituting *B. cretica* (199) with *B. oleracea* (175), which is not expected to be involved in the admixture event (*f4* = 0.00023, Z = 2.68). Two admixture events provide evidence of potential exoferal origins for at least two samples, *B. oleracea* (175) and *B. cretica* (199). The four-population tree of [[*B. montana* (222), curly kale],[*B. oleracea* (175), broccoli]] suggests significant gene flow from *B. montana* (222) to *B. oleracea* (175) (*f4* = 0.315, Z = 15.77), as does the tree of [[tronchuda cabbage, Chinese white kale],[*B. cretica* (199), broccoli] for gene flow from Chinese white kale to *B. cretica* (199) (*f4* = −0.009, Z = −7.98). The fifth added migration edge from *B. rupestris* to wild C - clade 2 explains the shared ancestry recovered in the fastSTUCTURE results. The test for treeness with [[curly kale, wild C - clade 2],[*B. rupestris, B. macrocarpa]]* indicated significant admixture from *B. rupestris* to Wild C - clade 2 (*f4* = −0.006, Z = −6.50), but was non-significant when substituting wild C - clade 2 with cauliflower (*f4* = −0.0003, Z = −0.338). Overall, these analyses highlight that the evolutionary history of *B. oleracea* is characterized by many admixture events and lineages of exoferal origins.

**Figure 4:**
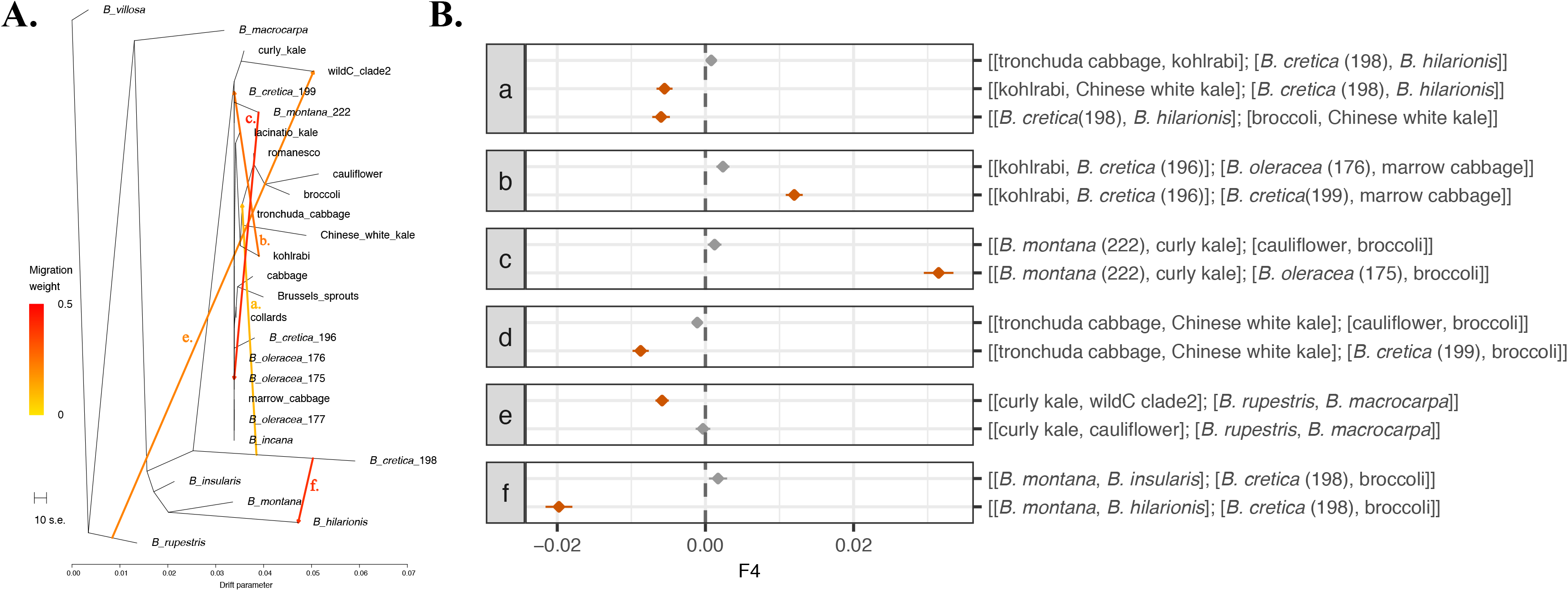
Inferred Admixture events. **A)** Phylogeny five migrations labeled a-f. **B)** Corresponding four-population tests for treeness.

### Archaeological and literary evidence point to a late-Holocene domestication

To further investigate the origins of domesticated *B. oleracea*, we surveyed archaeological, literary, and artistic evidence (***SI Appendix*, Table S5 & S6**). The earliest reported claim of *B. oleracea* comes from an archaeological collection from the Austrian Alps. This collection comprises three seeds dated to the Middle Bronze Age (ca. 3550-3350 years before present; BP (Schmidl and Oeggl 2005)). However, the lack of illustrations and discussion of separation criteria from other *Brassica* species makes us question the reliability of this species-level identification, as seeds of *Brassica* are difficult to tell apart. The only other find of similar antiquity is *B. oleracea* seeds from the Late Bronze Age/Early Iron Age, identified by scanning electron microscopy and radiocarbon dated directly between ca. 3250-2970 BP (Kaniewski et al. 2011). These finds are associated with destruction levels at Gibala, Tell Tweini in western Syria on the Mediterranean coast. While most of the archaeological finds are of seeds (***SI Appendix*, Table S5**), there is at least one documentation of pottery residues where lipids of *Brassica* leaf waxes were identified and dated to 850-750 BP (Evershed et al. 1992; Evershed et al. 1994). The authors attribute this to the boiling of leaves of *B. oleracea*, and given the lack of evidence for other commonly eaten *Brassica* leaves in England at this time, this would appear a likely identification.

The earliest literary references to *B. oleracea* date to Greek scholars 2500-2000 BP (***SI Appendix*, Table S6**). Hipponax’s writing refers to a seven-leaf cabbage in an iambic verse (West 2011), while Hippocrates *On the Nature of Women*, written around 2410-2320 BP, refers to the use of cabbage, or krambe, in a few recipes (Totelin 2009). As early as 2330 BP, there is evidence that the cultivar diversity that we are presently familiar with also existed historically. Eudermus refers to three kinds of cabbage: a smooth-leaved type, a curly-leaved type, and a salt variety, the latter he states has a delicate taste and appears to be distributed mainly in the eastern Aegean and on the modern Turkish coast (Yonge 1854). Theophrastus also refers to three varieties: a curly-leaved type, a smooth-leaved type, and a wild type with a bitter taste, many branches, and many small round leaves (Yonge 1854). Pliny in his *Natural History* writing some 200 years later describes at least an additional ten varieties than those seen in the previous classical works ((the Elder.) and Rackham 1950). However, while most scholars accept that the translations of Greek or Latin of cabbage refers to *B. oleracea*, it is important to note that both in the Greek-English Lexicon (Liddell and Scott 1940) and in Hort’s (Hort 1916) translation of Theophrastus’ *Historia Plantarum*, cabbage is translated as *B. cretica*, not *B. oleracea*. Certainly there are differences between the subspecies of *B. cretica* that might be reflective of such differences in the past and the diversity we see in our PCA plots. Further, the description by Niciander (quoted by Athenaeus; (Yonge 1854); p. 582) indicates that wild or at least feral forms of *B. cretica* were known in Ionia, the western coast of present day Turkey, ca. 2150-2050 BP.

### Late-Holocene environmental niche modeling highlights wild relatives’ ranges

Using information from archaeology and literature, we can infer that *B. oleracea* was under cultivation as early as 3550-3350 BP, around the late-Holocene. To predict what would be a suitable habitat for the wild relatives during the late-Holocene, we compiled occurrence records from GBIF (www.gbif.org) and (Snogerup et al. 1990), along with environmental data, to perform environmental niche modeling using MaxEnt 3.4.1 (Phillips et al. 2017). Notably, we find that *B. cretica* has an expanded Eastern Mediterranean habitat suitability (**Fig. 3C**) that includes Cyprus, where only *B. hiliarionis* is known presently. The only other species with a current day Eastern Mediterranean distribution is *B. hilarionis* (**Fig. 3B**), which also has expanded habitat suitability to the surrounding mainland coastal regions (**Fig. 3D**). However, since most of these wild species are narrow island endemics ((Snogerup et al. 1990), species are estimated to have little change from current day distributions (***SI Appendix*, Fig. S5 and Table S7**).

## DISCUSSION

### Multiple lines of evidence support a single Eastern Mediterranean origin

Our evidence from genome-scale, multilocus data along with archeology, literature, and environmental niche modeling best supports a single Eastern Mediterranean domestication origin for *B. oleracea*, corroborating the conclusions of (Maggioni et al. 2018) based on literary sources alone. When modeling phylogeny and population structure, two Eastern Mediterranean species, *B. cretica* and *B. hilarionis*, are found as sister species to cultivars and are assigned ancestry from all populations for values of K from 2 to 4 (**Fig. 1**), consistent with these species being likely parental species of *B. oleracea* domesticates. In our species tree reconstructions, we find just *B. cretica* as sister to all cultivars, specifically sample 198, which also clusters with wild-collected *B. cretica* samples from (Kioukis et al. 2020) in our PCA (Fig. 2A-B), lending further support for *B. cretica* as the progenitor species. This same sample of *B. cretica* (198) as well as our sample of *B. hilarionis* are recovered as fairly homozygous, therefore they would likely be good starting material for future research related to genome editing using wild relatives such as de novo domestication.

While we do recover evidence of admixture between *B. cretica* (198) and both wild and cultivated taxa, the placement of *B. cretica* (198) as the closest living wild relative does not change. However, an inferred admixture event from *B. cretica* (198) to *B. hilarionis* does result in a topological change in the placement of *B. hilarionis* as sister to *B. montana* (224; originally collected in Spain) (***SI Appendix*, Fig. S4**). This novel relationship has not been identified before and warrants additional study with greater taxon sampling. The second migration event involving *B. cretica* (198) is from Chinese white kale. This event lends further evidence of admixture with wild germplasm during the domestication process, consistent with other examples demonstrating that domestication is not a single event, but a series of events characterized by continuous gene flow between wild and cultivated populations (Beebe et al. 1997; Wang et al. 2017). Together with the phylogeographic discontinuity of wild *B. oleracea* samples and their Eastern Mediterranean progenitors (**Fig. 3B**), the more distant phylogenetic placement of *B. insularis, B. macrocarpa*, and *B. villosa* (**Fig. 3A**), and strong patterns of shared ancestry between *B. incana* and cultivars (**Fig. 1**), these results lead us to support the hypothesis of domestication in the Eastern Mediterranean with *B. cretica* as the closest living wild relative.

### The role of ferality in the domestication of *Brassica oleracea*

Multiple lines of evidence highlight the role of wild and feral populations as pools of diversity that contributed to crop diversification during domestication (Beebe et al. 1997; Allaby 2010; Fuller et al. 2014; Wang et al. 2017). Our data supports a similar phenomenon in the domestication of *B. oleracea*: it appears that introgression from wild or feral populations contributed to the genetic composition of particular crops, and vice versa, which is revealed by in-depth analyses of admixture using population structure and tree-based methods (**Fig. 1; Fig. 4; *SI Appendix*, Fig. S4**). Several wild relatives, including *B. cretica*, as well as wild *B. oleracea, B. incana*, and samples of *B. montana*, and *B. villosa*, are recovered as feral in all analyses.

While we find one sample of *B. cretica* (198) as the closest living wild relative, we also identify two samples of *B. cretica* (196 and 199; ***SI Appendix*, Fig. S6**) are likely feral and fall within the cultivar clade (**Fig. 1**). Interestingly, (Song et al. 1988) also recovered a polyphyletic *B. cretica* using RFLPs. Results presented here support previous findings that *B. cretica* was at one point domesticated. (Snogerup et al. 1990) state that wild *B. cretica* was consumed as late as 1962 and, as noted in our literary results, some early references to *B. oleracea* in the literature could be translated as *B. cretica*, meaning the vast amount of described morphology in these works, which may be the result of cultivation, could now be reflected in the multiple named subspecies and described genetic diversity of modern *B. cretica* (Snogerup et al. 1990; Widén et al. 2002; Allender et al. 2007; Edh et al. 2007). Further, the fact that feral forms of *B. cretica* were known in at least Ionia (western coast of present day Turkey) ca. 2150-2050 BP and the evidence of feral *B. cretica* populations today in Lebanon, which are morphologically similar to *B. cretica* subsp. *nivea*, suggests widespread trade of these species by the earliest Mediterranean civilizations (Dixon 2006). Previous researchers have also noted that *B. cretica* populations are typically found in coastal locations associated with ancient seaports, occupying their preferred ecological niche on chalk cliffs undisturbed by grazing (Mitchell 1976; Snogerup et al. 1990). We believe that these early forms of *B. cretica* may have played underappreciated roles in the domestication of *B. oleracea* crops and to fully understand the evolutionary history of *B. oleracea*, the domestication story of *B. cretica* must be resolved.

Sources have hypothesized wild populations of *B. oleracea* in England (Snogerup et al. 1990; Song et al. 1990) are the progenitor(s) for modern cultivars, while others have proposed that these are escaped cultivars (Mitchell 1976; Mitchell and Richards 1979). Consistent with these hypotheses, we find that the three wild *B. oleracea* samples in our study cluster with cultivars both phylogenetically and in PCA for both SNP data and expression profiles. Although these samples are from Canada (175), Denmark (176), and Germany (177) (***SI Appendix*, Fig. S6**), and notably do not include sampling of *B. oleracea* populations in England, one of the hypothesized geographic origins (**Table 1**), we suggest that an origin in England is unlikely given the archeological and literary data. Although the oldest archaeobotanical record for *B. oleracea* (Middle Bronze Age; ca. 3550 −3350 BP) is from Austria, we regard this evidence with caution as wild populations of *B. oleracea* are not presently found in Austria and the major *Brassica* crops in this region include *B. nigra* (Tutin 1964) or potentially cultivated turnip (*B. rapa*). Additionally, there is no compelling archaeological evidence to suggest the possible cultivation of cabbages in Europe prior to the Late Iron Age (2350-2050 BP) and Roman periods (1950-1650 BP), but there is evidence for knowledge of *B. oleracea* in Greece during this time ((Maggioni et al. 2018)***; SI Appendix*, Tables S5 and S6**). Overall, there are no records for *B. oleracea* from before this period within databases relating to the Eastern Mediterranean (Reihl 2014), Europe (Kroll 2001; Kroll 2005), Britain (Tomlinson and Hall 1996), the Czech Republic (Kreuz and Schäfer 2002), or within pre-dynastic and Pharaonic Egypt (Murray 2000), despite having documentation for other *Brassica* species. Evidence for *B. oleracea* in Europe does not start appearing until ca. 1850 BP, when the appearance of seeds increased and can be attributed to the spread of crops both within and on the periphery of the Roman Empire (Van der Veen 2011). Additionally, several studies that sampled wild *B. oleracea* populations in the British Isles (Mitchell 1976; Mitchell and Richards 1979), South West England (Raybould et al. 1999), Northern Spain (Gómez-Campo et al. 2005), Iberian Peninsula (Sánchez-Yélamo 2014), Atlantic coasts of western Europe (Mittell et al. 2020), and Atlantic coast of France (Maggioni et al. 2020) support that these wild *B. oleracea* populations are feral populations, typically with low levels of genetic diversity and some degree of isolation from other populations. (Lanner-Herrera et al. 1996) sampled populations across Spain, France, and Great Britain, concluding that each population evolved independently, while more recently (Mittell et al. 2020) found that geographically close populations were more genetically different than distant populations. Our results provide additional evidence that feralization is commonplace for *B. oleracea* crops and that references to wild *B. oleracea* likely represent multiple, independent feralization events. Additional sampling of wild populations will enable opportunities to further investigate the relationships among these feral populations and cultivated crops.

*Brassica incana*, another suggested progenitor species (Snogerup 1980), is also supported as feral by our analyses. Two of our five samples (204 and 207; ***SI Appendix*, Fig. S6**) are recovered as sister to all cultivars in our individual level phylogeny, but are found to share 100% of their ancestry with cultivars rather than other wild taxa using fastSTRUCTURE when K = 2 (**Fig. 1**). Further, these two samples were resolved as sister to lacinato kale in our species tree analysis, providing additional evidence that these samples represent a feral lineage, possibly of lacinato kale. This result may provide insight to why previous studies have found *B. incana* as sister to *B. oleracea (Lázaro and Aguinagalde 1998; Mei et al. 2010*) and the observation by (Snogerup et al. 1990) that some samples of *B. incana* are more interfertile with cultivated *B. oleracea* than others. Although (Snogerup et al. 1990) suggested that *B. incana* was more interfertile due to historical introgression, we do not find evidence for this for samples 204 and 207. However, the three other samples of *B. incana* (205, 208, 209), which are found belonging to the newly identified wild C - clade 2, do show evidence of admixture with *B. rupestris*. These three samples were collected in Italy, while the two other samples found in this clade, *B. cretica* (195) and *B. villosa* (233), were collected in Greece and Italy, respectively (***SI Appendix*, Fig. S7**). All five wild C - clade 2 samples share an introgression event from *B. rupestris* (**Fig. 1; Fig. 4; *SI Appendix*, Fig. S4**), but come from different germplasm collections (IPK-gatersleben and USDA National Plant Germplasm System), ruling out the inferred migration being the result of current cultivation practices. It is possible that these samples are related to the wild kale of Crimea, which is posited as a *B. rupestris–incana* hybrid which was transferred to the Crimea via trade (Dixon 2006). This suggests that there was early widespread cultivation of these *B. rupestris–incana* types (Dixon 2006) and provides a plausible explanation for why *B. incana* and *B. rupestris* are closely related in previous studies (Lannér et al. 1997; Mei et al. 2010).

The last feral identification is that of *B. montana*, for which we find one sample as more closely related to wild taxa (224) and one more closely related to cultivars (222) (***SI Appendix*, Fig. S6**). The feral sample (222) is of unknown origin, but again the literature indicates that this may not be a surprising result. Many studies have previously indicated a close relationship between *B. montana* and *B. oleracea*. For example, (Panda et al. 2003) concluded that *B. montana* may be a subspecies of *B. oleracea*, while (Lannér et al. 1997) found that *B. montana* and *B. oleracea* clustered together using chloroplast data. Furthermore, several authors have suggested that some populations of *B. montana* were feral *B. oleracea* (Paolucci 1890; Onno 1933; Snogerup et al. 1990), which may be reflected in the overlapping ranges produced by our niche modeling of these two species (***SI Appendix*, Fig. S5**). Therefore, in combination with results from previous studies, our results support that at least some *B. montana* populations are of feral origin.

Taken together, it is clear that the current taxonomy of *B. oleracea* and its wild relatives is confounded by gene flow between wild and cultivated populations, resulting in confusion between wild and feral lineages and obscuring the true evolutionary history of this species. Additionally, while there is much interest in crop improvement using CWRs (Meyer et al. 2012; Khoury et al. 2020), feral lineages offer another, potentially more direct route to reintroducing genetic diversity into cultivated populations, as gene flow is less likely to be impeded by barriers such as reproductive isolation (Mabry et al. 2021). These feral populations may also provide additional avenues to explore the evolutionary capacity for range expansion and phenotypic plasticity.

### Post-domestication cultivar relationships

While our knowledge of the spread and diversification of *B. oleracea* crops after domestication is compounded by both the difficulties of identifying seeds of individual crop types and frequent introgression between crop types, we can infer some patterns using the species phylogeny. Like other studies (Cheng et al. 2016; Stansell et al. 2018), we find Chinese white kale sister to all other cultivars, representing the only Asian clade of crop types (**Fig. 3A**). While the spread of *B. oleracea* to eastern Asia is still undocumented archaeologically, recent pollen analysis has provided evidence for cultivation of other *Brassica* species, including *B. rapa*, in the Yangtze valley 3250 - 3350 BP, likely corresponding to movement across “Silk Road” trade routes (Zhang 2009). However, this only provides identification criteria, not archaeological evidence (Yang et al. 2018). A review of Chinese historical sources concluded that *B. oleracea* may have been introduced to China 1450 −1350 BP and had evolved into Chinese white kale in Southern China by the period of the Tang Dynasty (1350 - 1250 BP; (Zhang 2009)). Due to both its position as sister to all other cultivars and as the only Asian *B. oleracea* crop type, this taxon warrants additional study to understand its own unique domestication story.

The dispersal of *B. oleracea* by human translocation westward, ultimately to the Atlantic coast of Europe, appears to have established both regional feral populations and the variety of modern crop types. Archaeological evidence suggests that this process may have begun with Late Bronze Age seafaring (3000-3300 years ago), when the whole Mediterranean became linked in trade perhaps for the first time (Broodbank 2015), and continued to provide a corridor for introgression and varietal diversification through the Iron Age (up to 2000 years ago). Trade links along the Atlantic seaboard from North Africa and Iberia through Britain and Ireland are clearly indicated in archaeology (Cunliffe 2004), and are associated with the first peopling of the Canary Islands from the north, where walking stick kale is endemic. Notably, many cultivars do not form monophyletic groups in our sample level phylogeny, likely indicative of admixture between crop types. This is supported by previous findings that broccoli is paraphyletic (Song et al. 1988; Stansell et al. 2018), as well as collards (Pelc et al. 2015), and by our findings that kale types such as tronchuda kale and perpetual kale are highly polyphyletic, suggesting that the kale-phenotype has been selected for multiple times independently.

In conclusion, we confirm a single Eastern Mediterranean origin for *B. oleracea* and find *B. cretica* as the closest living wild relative. We highlight several feral lineages that are not reflected by the current taxonomy but likely reflect important aspects of the domestication history for *B. oleracea*. Moving forward, it will be important to collect, study, and preserve these feral lineages as pools of allelic diversity, which may play an important role in future crop improvement, e.g. as a source of potential pest and pathogen resistance (Mithen et al. 1987; Mithen and Magrath 1992; Mohammed et al. 2010). In clarifying the evolutionary history of *B. oleracea* and its wild relatives, we hope to enable this model system for additional studies on evolutionary phenomena such as parallel selection, polyploidy, and ferality. Additionally, since many of these wild species are very narrow endemics and are valuable for both crop improvement and for nature conservation, their identification and preservation is urgent. We hope this study can serve as a stepping stone, as the work before us has, for all those who, like Darwin, are intrigued by this group of plants and wish to further its study.

## MATERIALS AND METHODS

### Taxon sampling

Samples from cultivars accounted for 188 of the 224 total samples with the remaining 36 samples included being previously identified wild relatives (***SI Appendix*, Table S2**). These include accessions from the United States Department of Agriculture, Agriculture Research Service (USDA-ARS) Plant Genetic Resources Unit (PGRU; 114 accessions), The Leibniz Institute of Plant Genetics and Crop Plant Research (IPK; 71 accessions), Universidad Politécnica de Madrid (UPM; 4 accessions), The Nordic Genetic Resource Centre (NordGen; 2 accessions), Gomez Campo Collection (2 accessions), John Innes Center (1 accession), doubled haploid lines (17 samples, some accessions sampled twice), or from the Pires’ personal collection (13 accessions). Four replicates of each accession were grown from seed in a sterile growth chamber at the University of Missouri (MU; Columbia, MO) Bond Life Sciences Center in a randomized complete block design across two independent outgrowths. At the seventh leaf stage, leaf four was collected from each plant and immediately flash-frozen in liquid nitrogen for RNA extraction. Morphotype identity was validated in mature plants by growing all accessions twice over the span of two years (***SI Appendix*, Table S2**).

Whole-genome resequencing data for an additional four samples from (Kioukis et al. 2020) of two varieties of *B. cretica* (var. *cretica* and var. *nivea*) was downloaded from the National Center for Biotechnology Information (NCBI) Sequence Read Archive (SRA) to supplement our sampling of *B. cretica*. These samples are under the SRA accession as follows: A = SRR9331103, B = SRR9331104, C = SRR9331105, and D = SRR9331106). Samples of A and B are *B. cretica* var. *nivea* from mainland Greece and C and D are *B. cretica* var. *cretica*, one from the mainland (C) and one from the island of Crete (D).

### RNA isolation and sequencing

RNA was isolated using the ThermoFisher Invitrogen PureLink RNA mini kit (Invitrogen, Carlsbad, CA, USA) followed by TruSeq library preparation (Illumina, San Diego, CA, USA) and sequencing on the NextSeq platform (Illumina, San Diego, CA, USA) for 2 × 75 bp reads. Library preparation and sequencing were performed through the MU DNA Core Facility. For eight flow cells, 24 samples were multiplexed and sequenced in a single flow-cell, followed by a ninth flow cell with 17 samples, and a tenth flow-cell with 16 samples.

### Mapping and SNP calling

Short reads were mapped to the *B. oleracea* TO1000 genome (Chinese white kale; (Parkin et al. 2014); release-41) by first using the STAR v. 2.5.2 (Dobin et al. 2013) two-pass alignment to identify splice junctions, which were then used in the second pass to improve mapping (Engström et al. 2013). The TO1000 genome of Chinese white kale was chosen due to wild relatives having a more kale-like phenotype and its placement as sister to the other cultivars in recent studies (Cheng et al. 2016; Stansell et al. 2018). Mapped reads (BAM format) were then processed following the GATK v. 3.8 best practices for RNA-seq reads (McKenna et al. 2010; Van der Auwera et al. 2013; Poplin et al. 2017). To ensure that reads were mapping correctly, the GATK ‘Split’N’Trim’ function was used to split reads into exon segments and trim any overhanging reads in intron segments. In total, 7,564,168 variants were called before any filtering was performed. The resulting variants were filtered to exclude those with a Fisher strand (FS) value greater than 30 and quality depth (QD) less than 2.0. This recovered 942,357 variants in total, with 879,865 variants on chromosomes 1-9 and 62,492 variants on scaffolds. Only variants aligning to chromosomes were used in downstream analyses. The remaining variants were then filtered using vcftools v. 0.1.17 (Danecek et al. 2011) to exclude sites with greater than 60% missing data (*--max-missing 0.4*), sites with mean depth values less than 5 (*--min-meanDP 5*), and indels (*--remove-indels;*) resulting in a total of 103,525 SNPs. Finally, SNPs were filtered for linkage disequilibrium (LD) using using PLINK v. 1.90 with a window size of 50 Kb, or about two times the estimated length for 80% LD decay (Cheng et al. 2016), a step size of 5 kb, and a variance inflation factor of 2 (*--indep 80kb 5 2;* (Purcell et al. 2007), for a final dataset of 30,014 SNPs). The four *B. cretica* genome resequencing samples (Kioukis et al. 2020) were also mapped to the *B. oleracea* TO1000 genome (Chinese white kale; (Parkin et al. 2014); release-41), using BWA (Li and Durbin 2009). For all samples no mapping bias was detected when comparing the percentage of uniquely mapped reads across cultivar groups, species, and sequencing lane (***SI Appendix*, Fig. S8**).

### Phylogenetic and Introgression Inference

To test how the different populations are related to one another and which wild relative is most closely related to the cultivated types, we used three different phylogenetic programs; SNPhylo v. 20160204 (Lee et al. 2014) to assess individual sample relationships, IQ-Tree v. 1.6 (Nguyen et al. 2015) to test species level relationships, and TreeMix v. 1.13 (Pickrell and Pritchard 2012) to assess introgression. For SNPhylo (Lee et al. 2014), `we ran analyses using 0.1 for LD, minor allele frequency ≥ 0.01, proportion of missing sites ≤ 0.4, 1000 bootstrap replicates, and rooted with sample 238` (*B. villosa*). For IQ-Tree, we used the Polymorphism-aware phylogenetic Models (PoMo) software (Schrempf et al. 2016; -*m GTR+P*) to perform phylogenetic comparisons using population genetic data, using 1000 bootstrap replicates via the ultrafast bootstrap approximation method (Hoang et al. 2018) and *B. villosa* to root the tree. For our IQ-Tree analysis, we subsampled data to include only those samples which were recovered as monophyletic in our SNPhylo tree (***SI Appendix*, Table S2; sample # with asterisks).** To test both the topology of relationships and for gene flow between populations, we used TreeMix with the following parameters: no sample size correction (-*noss*), rooted with *B. villosa (-root villosa*), bootstrapping over blocks of 500 SNPs (-*bootstrap -k 500*), and to incorporate between 2-10 migration events (-*m*). TreeMix (Pickrell and Pritchard 2012) was run with samples of *B. cretica, B. incana, B. montana*, and *B. oleracea* as individuals, but used samples found in wild C- clade 2, cultivars, and wild relatives as populations. Four-population (*f4*) tests for treeness (Reich et al. 2009; Pickrell and Pritchard 2012) were used to test the support of the inferred migration edges from Treemix (Pickrell and Pritchard 2012) via the fourpop method.

### Population Structure and Variation

To test ancestry proportions and identify the likely genetic structure of described populations we used fastSTRUCTURE v. 1.0 (Raj et al. 2014). We tested K values from 2 to 4 using default convergence criteria and priors followed by the *chooseK.py* script to determine the appropriate number of model components that best explain structure in the dataset.

Angsd v. 0.925 (Korneliussen et al. 2014) was used to calculate genotype likelihoods for all samples, plus the four additional *B. cretica* samples from (Kioukis et al. 2020), using the parameters -*doGlf 2 - doMajorMinor 1 -doMaf 2 -minMapQ 30 -SNP_pval 1e-6*, followed by analysis with PCAangsd v. 0.97 (Meisner and Albrechtsen 2018) to visualize population structure, estimate allele frequencies, and calculate individual inbreeding coefficients using the parameters -*admix -selection 1 -inbreed 2*.

### Clustering based on Expression Profiles

First, Salmon v. 1.2.1 (Patro et al. 2015) was used to acquire transcript abundances for each sample and the estimated number of reads originating from transcripts. The input for expression profile analysis was prepared using tximport (Soneson et al. 2015) with design = ~plantout + cultivar type. Correction for library size (*estimateSizeFactors*) and variance-stabilizing transformation (*vst*) was performed in DESeq2 v. 1.28.1 (Love et al. 2014). To test for clustering based on expression profiles, we ran a PCA on the normalized expression values and performed clustering based on Euclidean distance using the ‘prcomp’ and ‘hclust’ functions, respectively, in the ‘stats’ v. 3.6.2 package for R v. 3.6.0 (R Core Team 2018). To assess networks of genes driving differences observed in the PCA, we used WGCNA v. 1.68 (Langfelder and Horvath 2008). Following (Zhang and Horvath 2005), we found that a soft-thresholding power of nine was best as it was the lowest power that satisfied the approximate scale-free topology criterion, resulting in 48 modules of genes.

To determine biological processes which were overrepresented in the resulting modules, *Arabidopsis thaliana* orthologs of *B. oleracea* were determined using both synteny and BLAST. Synteny-based annotations were extracted from Table S7 in (Parkin et al. 2014) while the BLAST annotation was performed using blastn in BLAST v. 2.10.0+ (Camacho et al. 2009). The *B. oleracea* CDS database was downloaded from https://plants.ensembl.org/Brassica_oleracea/Info/Index, and the *A. thaliana* CDS database from Araport11_genes.201606.cds.fasta from https://www.arabidopsis.org/. The blastn parameters were -*evalue 1E-6 -max_target_seqs 1*. Genes determined using synteny were then used to perform a GO analysis through PANTHER v. 16.0 (Mi et al. 2021).

### Environmental niche modeling

We compiled occurrence records for wild relatives from the Global Biodiversity Information Facility (GBIF, www.gbif.org) data portal and data from (Snogerup et al. 1990). From the GBIF data, we omitted records that were duplicated, lacked location data and/or vouchers, were collected from the grounds of botanical gardens, and that were clearly outside of the native range. From the (Snogerup et al. 1990) data, we omitted records that could not be georeferenced to <5km spatial uncertainty. Populations of *B. cretica* in Lebanon and Israel and of *B. incana* in Crimea are thought to be likely early human introductions (Snogerup et al. 1990) and records from these areas were omitted. Occurrences above 1200m altitude were also omitted, as these species rarely occur above 1000m and observations above these altitudes may represent anthropogenic dispersals to disturbed areas or misidentifications. To minimize sampling bias due to clustered observations (Beck et al. 2014; Boria et al. 2014), we thinned the filtered occurrences to records greater than or equal to 10km apart using the ‘spThin’ package in R (Aiello-Lammens et al. 2015). After filtering and thinning, 172 records remained for *B. cretica*, 65 for *B. incana*, 57 for *B. insularis*, 101 *B. montana*, 15 for *B. villosa*, and 7 and 6 for the narrow endemics *B. macrocarpa* and *B. hilarionis* respectively. Next, we obtained rasters for 19 bioclimatic variables at 2.5 minutes resolution based on contemporary climate data from WorldClim v. 2.0 (Fick and Hijmans 2017) and rasters for 19 bioclimatic variables at 2.5 minutes resolution based on late-Holocene climate projections using data derived from PaleoClim (Fordham et al. 2017; Brown et al. 2018). Rasters were clipped using QGIS v. 3.83 (Open Source Geospatial Foundation Project) to constrain the geographical background to windows slightly larger than the area circumscribed by contemporary observational data (Phillips et al. 2009; Acevedo et al. 2012). While it is common practice to eliminate collinear environmental variables to avoid overfitting (Braunisch et al. 2013), recent simulations have shown that removing highly collinear variables has an insignificant impact on maximum entropy model performance (Feng et al. 2019) so all original variables were included. Projections for late-Holocene habitat suitability were generated using MaxEnt v. 3.4.1 (Phillips et al. 2017). Linear, quadratic, product, and hinge features and jackknife resampling was used to measure variable importance. Relative model performance was evaluated with the adjusted area under receiver operating characteristic (ROC) curve (AUC; (DeLong et al. 1988)). While optimal performance cannot be determined with this approach using presence-only data, relative performance can still be assessed (Phillips et al. 2006).

## DATA AVAILABILITY

The sequences reported in this paper have been deposited in the Sequence Read Archive database (accession no. PRJNA544934).

## AUTHOR CONTRIBUTIONS

MEM, SDTH, ACM, HA, PPE, JDM, DACP, GRT, CJS, GB, JL, DQF, TB, RGA, JED, MAG, and JCP designed the project. MEM, EYG, HA, and SDTH grew plants and collected tissue. MEM and EYG extracted and isolated RNA. MEM analyzed the genetic data. ACM produced the species distribution models. CS, DQF, and RGA researched archeology and written data. SDTH, HA, and JED assisted with processing and analyzing the data. MEM wrote the original manuscript. PPE, JDM, DACP, GRT, CJS, GB, JL, TB, MAG, and JCP provided critical feedback on manuscript drafts.

## ACKNOWLEDGMENTS

We thank Drs. Bob Schnabel, Troy Rowan, Harly Durbin, and Paul Blischak for their assistance with computational analyses, the Mizzou DNA core, Nathan Bivens, Ming-Yi Zhou, and Karen Bromert, for their assistance in getting quality data for sequencing, and our computing resources, specifically the Research Computing Support Services (RCSS) and Informatics Research Core Facility (IRCF) at the University of Missouri. We thank Sarah Unruh for valuable feedback on early versions of this manuscript and Dr. Jeff Ross-Ibarra for his help with interpreting admixture statistics. Finally, we thank our funding USDA-ARS Project No. 8060-21000-024-00D and the National Science Foundation Postdoctoral Fellowship in Biology (Award No. 1711347, S.T-H).

## TABLE LEGENDS

**Table 1.** Wild species which have been proposed as progenitor species for *B. oleracea* crop types. Specific location is included in parentheses if indicated by the author.

## SUPPLEMENTARY MATERIAL

**Table S1:** *Brassica oleracea* crop types with common name, species name, Kew cultivar group, and other used names. Illustrations by Andi Kur.

**Table S2:** Sample information with species, variety, cultivar, accession/collection, sample #, and SRA #. Asterisks (*) next to sample # indicate those samples that were recovered as monophyletic and used in species tree reconstruction.

**Table S3:** WGCNA predicted gene modules with number of genes in each module, the number of annotated *Arabidopsis thaliana* genes using blast and synteny, and the percent of syntenic genes represented in the module. Module number with asterisks (*) represent the five modules with the largest percent of syntenic genes in the module.

**Table S4:** Top five WGCNA modules with largest percent of syntenic genes represented in the module with corresponding annotated GO biological process with the largest fold enrichment, and p-value.

**Table S5:** Archaeological *Brassica* reports from Europe and the Eastern Mediterranean.

**Table S6:** Literary and artistic sources covering the Classical Greek, Roman, and medieval and post-medieval sources.

**Table S7:** Area under the receiver operating characteristic curve (AUC) values for Maxent environmental niche model runs for putative wild relatives of Brassica oleracea crops.

**Figure S1:** PCAs of SNPs and Expression profiles. **A)** Genetic variation PCA of PC3 vs PC4, **B)** Expression profile PCA for PC2 vs PC3, and **C)** Expression profile PCA for PC3 vs PC4 of wild and cultivar samples. Triangles = wild samples, circles = cultivars. Triangles with black outlines = wild C - clade 2 samples with species identification indicated by color. Wild-collected *B. cretica* samples from (Kioukis et al. 2020) indicated by asterisks, labeled as SRA.

**Figure S2:** Inbreeding coefficients for wild, cultivar, and feral samples.

**Figure S3:** *Brassica oleracea* cultivars and wild relative dendrogram based on expression profiles. Wild species indicated by color below. Cultivars in grey below. Wild C - clade 2 indicated by black outlines for the corresponding bar chart below. Sample names in red = putatively wild samples.

**Figure S4:** TreeMix analysis of wild, cultivar, and feral samples. **A-F)** Phylogeny with 0-5 migrations indicated.

**Figure S5:** The current wild relatives distribution and modeled late-Holocene suitable habitat. Middle - current species distribution of wild relatives. Suitable habitat for **A)** *B. montana*, **B)** *B. insularis*, **C)** *B. macrocarpa*, **D)** *B. rupestris*, **E)** *B. villosa*, **F)** *B. incana*, and **G)** *B. oleracea*, during the late-Holocene

**Figure S6:** Leaf scans of feral samples. Leaf used for RNA collection with biological replicate indicated.

**Figure S7:** Leaf scans of wild C - clade 2 samples. Leaf used for RNA collection with biological replicate indicated.

**Figure S8:** Mapping percentage of unique reads for **A)** wild and cultivated samples, **B)** cultivar groups, and **C)** sequencing lane.

## REFERENCES

Acevedo P, Jiménez-Valverde A, Lobo JM, Real R. 2012. Delimiting the geographical background in species distribution modelling. Journal of Biogeography [Internet] 39:1383–1390. Available from: http://dx.doi.org/10.1111/j.1365-2699.2012.02713.x

Aiello-Lammens ME, Boria RA, Radosavljevic A, Vilela B, Anderson RP. 2015. spThin: an R package for spatial thinning of species occurrence records for use in ecological niche models. Ecography [Internet] 38:541–545. Available from: http://dx.doi.org/10.1111/ecog.01132

Allaby R. 2010. Integrating the processes in the evolutionary system of domestication. Journal of Experimental Botany [Internet] 61:935–944. Available from: http://dx.doi.org/10.1093/jxb/erp382

Allender CJ, Allainguillaume J, Lynn J, King GJ. 2007. Simple sequence repeats reveal uneven distribution of genetic diversity in chloroplast genomes of Brassica oleracea L. and (n = 9) wild relatives. Theor. Appl. Genet. 114:609–618.

Bailey LH. 1930. The Cultivated Brassicas Second Paper. Gentes Herbarum.

Beck J, Böller M, Erhardt A, Schwanghart W. 2014. Spatial bias in the GBIF database and its effect on modeling species’ geographic distributions. Ecol. Inform. 19:10–15.

Beebe S, Toro Ch O, Gonza’lez AV, Chaco’n MI, Debouck DG. 1997. Wild-weed-crop complexes of common bean (Phaseolus vulgaris L., Fabaceae) in the Andes of Peru and Colombia, and their implications for conservation and breeding. Genet. Resour. Crop Evol. 44:73–91.

Boria RA, Olson LE, Goodman SM, Anderson RP. 2014. Spatial filtering to reduce sampling bias can improve the performance of ecological niche models. Ecol. Modell. 275:73–77.

Braunisch V, Coppes J, Arlettaz R, Suchant R, Schmid H, Bollmann K. 2013. Selecting from correlated climate variables: a major source of uncertainty for predicting species distributions under climate change. Ecography 36:971–983.

Broodbank C. 2015. The Making of the Middle Sea: A History of the Mediterranean from the Beginning to the Emergence of the Classical World. Thames & Hudson

Brown JL, Hill DJ, Dolan AM, Carnaval AC, Haywood AM. 2018. PaleoClim, high spatial resolution paleoclimate surfaces for global land areas. Sci Data 5:180254.

Camacho C, Coulouris G, Avagyan V, Ma N, Papadopoulos J, Bealer K, Madden TL. 2009. BLAST+: architecture and applications. BMC Bioinformatics 10:421.

de Candolle A. 1855. Géographie botanique raisonnée ou exposition des faits principaux et des lois concernant la distribution géographique des plantes de l’époque actuelle. V. Masson

Cheng F, Sun R, Hou X, Zheng H, Zhang F, Zhang Y, Liu B, Liang J, Zhuang M, Liu Y, et al. 2016. Subgenome parallel selection is associated with morphotype diversification and convergent crop domestication in Brassica rapa and Brassica oleracea. Nat. Genet. 48:1218–1224.

Cunliffe B. 2004. Facing the Ocean: The Atlantic and Its Peoples, 8000 BC-AD 1500. Oxford University Press

Danecek P, Auton A, Abecasis G, Albers CA, Banks E, DePristo MA, Handsaker RE, Lunter G, Marth GT, Sherry ST, et al. 2011. The variant call format and VCFtools. Bioinformatics [Internet] 27:2156–2158. Available from: http://dx.doi.org/10.1093/bioinformatics/btr330

Darwin C, Gray A. 1868. The variation of animals and plants under domestication / by Charles Darwin; authorized edition, with a preface by Asa Gray. Available from: http://dx.doi.org/10.5962/bhl.title.37659

DeLong ER, DeLong DM, Clarke-Pearson DL. 1988. Comparing the areas under two or more correlated receiver operating characteristic curves: a nonparametric approach. Biometrics 44:837–845.

Dixon GR. 2006. Origins and diversity of Brassica and its relatives. Vegetable brassicas and related crucifers: 1–33.

Dobin A, Davis CA, Schlesinger F, Drenkow J, Zaleski C, Jha S, Batut P, Chaisson M, Gingeras TR. 2013. STAR: ultrafast universal RNA-seq aligner. Bioinformatics 29:15–21.

Edh K, Widén B, Ceplitis A. 2007. Nuclear and chloroplast microsatellites reveal extreme population differentiation and limited gene flow in the Aegean endemic Brassica cretica (Brassicaceae). Mol. Ecol. 16:4972–4983.

Engström PG, Steijger T, Sipos B, Grant GR, Kahles A, Rätsch G, Goldman N, Hubbard TJ, Harrow J, Guigó R, et al. 2013. Systematic evaluation of spliced alignment programs for RNA-seq data. Nat. Methods 10:1185–1191.

Evershed RP, Arnot KI, Collister J, Eglinton G, Charters S. 1994. Application of isotope ratio monitoring gas chromatography–mass spectrometry to the analysis of organic residues of archaeological origin. The Analyst [Internet] 119:909–914. Available from: http://dx.doi.org/10.1039/an9941900909

Evershed RP, Heron C, Charters S, Goad LJ. 1992. The survival of food residues: new methods of analysis, interpretation and application. In: Proceedings of the British Academy. Vol. 77. p. 2.

Farnham MW. 1996. Genetic variation among and within United States collard cultivars and landraces as determined by randomly amplified polymorphic DNA markers. J. Am. Soc. Hortic. Sci. 121:374–379.

Feng X, Park DS, Liang Y, Pandey R, Papeş M. 2019. Collinearity in ecological niche modeling: Confusions and challenges. Ecology and Evolution [Internet] 9:10365–10376. Available from: http://dx.doi.org/10.1002/ece3.5555

Fernie AR, Yan J. 2019. De Novo Domestication: An Alternative Route toward New Crops for the Future. Mol. Plant 12:615–631.

Fick SE, Hijmans RJ. 2017. WorldClim 2: new 1-km spatial resolution climate surfaces for global land areas. Int. J. Climatol. [Internet]. Available from: https://rmets.onlinelibrary.wiley.com/doi/abs/10.1002/joc.5086

Fordham DA, Saltré F, Haythorne S, Wigley TML, Otto-Bliesner BL, Chan KC, Brook BW. 2017. PaleoView: a tool for generating continuous climate projections spanning the last 21 000 years at regional and global scales. Ecography 40:1348–1358.

Fuller DQ, Denham T, Arroyo-Kalin M, Lucas L, Stevens CJ, Qin L, Allaby RG, Purugganan MD. 2014. Convergent evolution and parallelism in plant domestication revealed by an expanding archaeological record. Proceedings of the National Academy of Sciences [Internet] 111:6147–6152. Available from: http://dx.doi.org/10.1073/pnas.1308937110

Gómez-Campo C, Aguinagalde I, Ceresuela JL, Lázaro A, Martínez-Laborde JB, Parra-Quijano M, Simonetti E, Torres E, Tortosa ME. 2005. An exploration of wild Brassica oleracea L. germplasm in Northern Spain. Genet. Resour. Crop Evol. 52:7–13.

Gustafsson M, Bentzer B, Von Bothmer B, Snogerup. 1976. Meiosis in Greek Brassica of the oleracea group. Available from: https://pascal-francis.inist.fr/vibad/index.php?action=getRecordDetail&idt=PASCAL7637010832

Heaney RK, Roger Fenwick G, Mithen RF, Lewis BG. 1987. Glucosinolates of wild and cultivated Brassica species. Phytochemistry [Internet] 26:1969–1973. Available from: http://dx.doi.org/10.1016/s0031-9422(00)81740-9

Helm J. 1963. Morphologisch-taxonomische Gliederung der Kultursippen vonBrassica oleracea L. Die Kulturpflanze [Internet] 11:92–210. Available from: http://dx.doi.org/10.1007/bf02136113

Hoang DT, Chernomor O, von Haeseler A, Minh BQ, Vinh LS. 2018. UFBoot2: Improving the Ultrafast Bootstrap Approximation. Molecular Biology and Evolution [Internet] 35:518–522. Available from: http://dx.doi.org/10.1093/molbev/msx281

Hodgkin T. 1995. Cabbages, kales, etc. Brassica oleracea:76–82.

Hort A. 1916. Theophrastus: enquiry into plants. Cambridge (Mass.) and.

Kaniewski D, Van Campo E, Van Lerberghe K, Boiy T, Vansteenhuyse K, Jans G, Nys K, Weiss H, Morhange C, Otto T, et al. 2011. The Sea Peoples, from cuneiform tablets to carbon dating. PLoS One 6:e20232.

Khoury CK, Carver D, Greene SL, Williams KA, Achicanoy HA, Schori M, León B, Wiersema JH, Frances A. 2020. Crop wild relatives of the United States require urgent conservation action. Proc. Natl. Acad. Sci. U. S. A. 117:33351–33357.

Kioukis A, Michalopoulou VA, Briers L, Pirintsos S, Studholme DJ, Pavlidis P, Sarris PF. 2020. Intraspecific diversification of the crop wild relative Brassica cretica Lam. using demographic model selection. BMC Genomics 21:48.

Korneliussen TS, Albrechtsen A, Nielsen R. 2014. ANGSD: Analysis of Next Generation Sequencing Data. BMC Bioinformatics 15:356.

Kreuz A, Schäfer E. 2002. A new archaeobotanical database program. Veg. Hist. Archaeobot. 11:177–180.

Kroll H. 2001. Literature on archaeological remains of cultivated plants (1999/2000). Vegetation History and Archaeobotany [Internet] 10:33–60. Available from: http://dx.doi.org/10.1007/pl00013368

Kroll H. 2005. Literature on archaeological remains of cultivated plants 1981--2004. Available at archaeobotany. de/database. html. Accessed January 2:2016.

Langfelder P, Horvath S. 2008. WGCNA: an R package for weighted correlation network analysis. BMC Bioinformatics 9:559.

Lannér C, Bryngelsson T, Gustafsson M. 1997. Relationships of wild Brassica species with chromosome number 2 n= 18, based on RFLP studies. Genome 40:302–308.

Lanner-Herrera C, Gustafeson M, Filt AS, Bryngelsson T. 1996. Diversity in natural populations of wild Brassica oleracea as estimated by isozyme and RAPD analysis. Genet. Resour. Crop Evol. 43:13–23.

Lázaro A, Aguinagalde I. 1998. Genetic Diversity in Brassica oleracea L. (Cruciferae) and Wild Relatives (2 n= 18) using Isozymes. Ann. Bot. 82:821–828.

Lee T-H, Guo H, Wang X, Kim C, Paterson AH. 2014. SNPhylo: a pipeline to construct a phylogenetic tree from huge SNP data. BMC Genomics 15:162.

Liddell HG, Scott R. 1940. A Greek-English Lexicon Perseus.

Li H, Durbin R. 2009. Fast and accurate short read alignment with Burrows-Wheeler transform. Bioinformatics [Internet] 25:1754–1760. Available from: http://dx.doi.org/10.1093/bioinformatics/btp324

Li T, Yang X, Yu Y, Si X, Zhai X, Zhang H, Dong W, Gao C, Xu C. 2018. Domestication of wild tomato is accelerated by genome editing. Nature Biotechnology [Internet] 36:1160–1163. Available from: http://dx.doi.org/10.1038/nbt.4273

Lizgunova TV. 1959. The history of botanical studies of the cabbage, Brassica oleracea L. Bulletin of Applied Botany, Genetics and Plant Breeding 32:37–70.

Love MI, Huber W, Anders S. 2014. Moderated estimation of fold change and dispersion for RNA-seq data with DESeq2. Genome Biol. 15:550.

Mabry ME, Rowan TN, Pires JC, Decker JE. 2021. Feralization: Confronting the Complexity of Domestication and Evolution. Trends Genet. [Internet]. Available from: http://dx.doi.org/10.1016/j.tig.2021.01.005

Maggioni L, von Bothmer R, Poulsen G, Aloisi KH. 2020. Survey and genetic diversity of wild Brassica oleracea L. germplasm on the Atlantic coast of France. Genetic Resources and Crop Evolution [Internet] 67:1853–1866. Available from: http://dx.doi.org/10.1007/s10722-020-00945-0

Maggioni L, von Bothmer R, Poulsen G, Lipman E. 2018. Domestication, diversity and use of Brassica oleracea L., based on ancient Greek and Latin texts. Genetic Resources and Crop Evolution [Internet] 65:137–159. Available from: http://dx.doi.org/10.1007/s10722-017-0516-2

McKenna A, Hanna M, Banks E, Sivachenko A, Cibulskis K, Kernytsky A, Garimella K, Altshuler D, Gabriel S, Daly M, et al. 2010. The Genome Analysis Toolkit: a MapReduce framework for analyzing next-generation DNA sequencing data. Genome Res. 20:1297–1303.

Mei J, Li Q, Yang X, Qian L, Liu L, Yin J, Frauen M, Li J, Qian W. 2010. Genomic relationships between wild and cultivated Brassica oleracea L. with emphasis on the origination of cultivated crops. Genet. Resour. Crop Evol. 57:687–692.

Meisner J, Albrechtsen A. 2018. Inferring Population Structure and Admixture Proportions in Low-Depth NGS Data. Genetics 210:719–731.

Meyer RS, DuVal AE, Jensen HR. 2012. Patterns and processes in crop domestication: an historical review and quantitative analysis of 203 global food crops. New Phytol. 196:29–48.

Mi H, Ebert D, Muruganujan A, Mills C, Albou L-P, Mushayamaha T, Thomas PD. 2021. PANTHER version 16: a revised family classification, tree-based classification tool, enhancer regions and extensive API. Nucleic Acids Res. 49:D394–D403.

Mitchell ND. 1976. The status of Brassica oleracea L. subsp. oleracea (wild cabbage) in the British Isles. Watsonia 11:97–103.

Mitchell ND, Richards AJ. 1979. Brassica Oleracea L. ssp. Oleracea (B. sylvestris (L.) Miller). J. Ecol. 67:1087–1096.

Mithen RF, Lewis BG, Heaney RK, Fenwick GR. 1987. Resistance of leaves of Brassica species to Leptosphaeria maculans. Trans. Br. Mycol. Soc. 88:525–531.

Mithen RF, Magrath R. 1992. Glucosinolates and Resistance to Leptosphaeria maculans in Wild and Cultivated Brassica Species. Plant Breeding [Internet] 108:60–68. Available from: http://dx.doi.org/10.1111/j.1439-0523.1992.tb00100.x

Mittell EA, Cobbold CA, Ijaz UZ, Kilbride EA, Moore KA, Mable BK. 2020. Feral populations of Brassica oleracea along Atlantic coasts in western Europe. Ecol. Evol. 10:11810–11825.

Mohammed A, Addo-Quaye A, Asare-bediako E. 2010. Control of Diamond back Moth (Plutella xylostella) on Cabbage (Brassica oleracea var capitata) using Intercropping with Non-Host Crops “E. Asare-Bediako,” AA Addo-Quaye and “A. Mohammed” Department of Crop Science, University of Cape Coast, Cape Coast, Ghana. Am. J. Food Technol. 5:269–274.

Murray MA. 2000. Fruits, vegetables, pulses and condiments. In: Cambridge University Press.

Neutrofal F. 1927. Zytologische Studien über die Kulturrassen von Brassica oleracea. Oesterr. Bot. Z. 76:105–115.

Nguyen L-T, Schmidt HA, von Haeseler A, Minh BQ. 2015. IQ-TREE: a fast and effective stochastic algorithm for estimating maximum-likelihood phylogenies. Mol. Biol. Evol. 32:268–274.

Onno M. 1933. Die Wildformen aus dem Verwandtschaftskreis “Brassica oleracea L.”. Österreichische Botanische Zeitschrift 82:309–334.

Open Source Geospatial Foundation Project. QGIS Geographic Information System. QGIS.org [Internet]. Available from: http://qgis.org

Panda S, Martín J, Aguinagalde I. 2003. Chloroplast and nuclear DNA studies in a few members of the Brassica oleracea L. group using PCR-RFLP and ISSR-PCR markers: a population genetic analysis. Theor. Appl. Genet. 106:1122–1128.

Paolucci L. 1890. Flora marchigiana. Stab. tipo-lit. Federici

Parkin IAP, Koh C, Tang H, Robinson SJ, Kagale S, Clarke WE, Town CD, Nixon J, Krishnakumar V, Bidwell SL, et al. 2014. Transcriptome and methylome profiling reveals relics of genome dominance in the mesopolyploid Brassica oleracea. Genome Biol. 15:R77.

Patro R, Duggal G, Kingsford C. 2015. Salmon: Accurate, Versatile and Ultrafast Quantification from RNA-seq Data using Lightweight-Alignment. Cold Spring Harbor Laboratory [Internet]:021592. Available from: https://www.biorxiv.org/content/10.1101/021592v1.abstract

Pelc SE, Couillard DM, Stansell ZJ, Farnham MW. 2015. Genetic Diversity and Population Structure of Collard Landraces and their Relationship to Other Brassica oleracea Crops. Plant Genome 8:eplantgenome2015.04.0023.

Phillips SJ, Anderson RP, Dudík M, Schapire RE, Blair ME. 2017. Opening the black box: an open-source release of Maxent. Ecography [Internet] 40:887–893. Available from: http://dx.doi.org/10.1111/ecog.03049

Phillips SJ, Anderson RP, Schapire RE. 2006. Maximum entropy modeling of species geographic distributions. Ecol. Modell. 190:231–259.

Phillips SJ, Dudík M, Elith J, Graham CH, Lehmann A, Leathwick J, Ferrier S. 2009. Sample selection bias and presence-only distribution models: implications for background and pseudo-absence data. Ecol. Appl. 19:181–197.

Pickrell JK, Pritchard JK. 2012. Inference of population splits and mixtures from genome-wide allele frequency data. PLoS Genet. 8:e1002967.

Poplin R, Ruano-Rubio V, DePristo MA, Fennell TJ, Carneiro MO, Van der Auwera GA, Kling DE, Gauthier LD, Levy-Moonshine A, Roazen D, et al. 2017. Scaling accurate genetic variant discovery to tens of thousands of samples. Cold Spring Harbor Laboratory [Internet]:201178. Available from: https://www.biorxiv.org/content/10.1101/201178v2.abstract

Prohens J, Gramazio P, Plazas M, Dempewolf H, Kilian B, Díez MJ, Fita A, Herraiz FJ, Rodríguez-Burruezo A, Soler S, et al. 2017. Introgressiomics: a new approach for using crop wild relatives in breeding for adaptation to climate change. Euphytica [Internet] 213. Available from: http://dx.doi.org/10.1007/s10681-017-1938-9

Purcell S, Neale B, Todd-Brown K, Thomas L, Ferreira MAR, Bender D, Maller J, Sklar P, de Bakker PIW, Daly MJ, et al. 2007. PLINK: a tool set for whole-genome association and population-based linkage analyses. Am. J. Hum. Genet. 81:559–575.

Raj A, Stephens M, Pritchard JK. 2014. fastSTRUCTURE: variational inference of population structure in large SNP data sets. Genetics 197:573–589.

Raybould AF, Mogg RJ, Clarke RT, Gliddon CJ, Gray AJ. 1999. Variation and population structure at microsatellite and isozyme loci in wild cabbage (Brassica oleracea L.) in Dorset (UK). Genet. Resour. Crop Evol. 46:351–360.

R Core Team. 2018. R: A language and environment for statistical computing. Available from: https://www.R-project.org/.

Reich D, Thangaraj K, Patterson N, Price AL, Singh L. 2009. Reconstructing Indian population history. Nature 461:489–494.

Reihl S. 2014. Archaeobotanical database of Eastern Mediterranean and Near Eastern sites. Available from: http://www.ademnes.de/

Sánchez-Yélamo MD. 2014. Characterisation of wild cabbage (Brassica oleracea L.) based on isoenzyme data. Considerations on the current status of this taxon in Spain. Genet. Resour. Crop Evol. 61:1295–1306.

Schiemann E. 1932. Entstehung der kulturpflanzen. Borntraeger

Schmidl A, Oeggl K. 2005. Subsistence strategies of two Bronze Age hill-top settlements in the eastern Alps—Friaga/Bartholomäberg (Vorarlberg, Austria) and Ganglegg/Schluderns (South Tyrol, Italy). Veg. Hist. Archaeobot. 14:303–312.

Schrempf D, Minh BQ, De Maio N, von Haeseler A, Kosiol C. 2016. Reversible polymorphism-aware phylogenetic models and their application to tree inference. J. Theor. Biol. 407:362–370.

Schulz OE. 1936. Cruciferae. In: Engler A, Harms H, editors. Die natuÈrlichen P anzenfamilien, 2nd edn. p. 176: 227±658.

Shyam P, Wu XM, Bhat SR. 2012. History, evolution, and domestication of Brassica crops. Plant Breed. Rev. [Internet]. Available from: https://www.cabdirect.org/cabdirect/abstract/20123077074

Snogerup S. 1980. The wild forms of the Brassica oleracea group (2n= 18) and their possible relations to the cultivated ones. Brassica crops and wild allies. [I].: 121–132.

Snogerup S, Gustafsson M, Von Bothmer R. 1990. Brassica sect. Brassica (Brassicaceae) I. Taxonomy and Variation. Willdenowia 19:271–365.

Soneson C, Love MI, Robinson MD. 2015. Differential analyses for RNA-seq: transcript-level estimates improve gene-level inferences. F1000Res. 4:1521.

Song KM, Osborn TC, Williams PH. 1988. Brassica taxonomy based on nuclear restriction fragment length polymorphisms (RFLPs). Theor. Appl. Genet. 75:784–794.

Song K, Osborn TC, Williams PH. 1990. Brassica taxonomy based on nuclear restriction fragment length polymorphisms (RFLPs). Theor. Appl. Genet. 79:497–506.

Stansell Z, Hyma K, Fresnedo-Ramírez J, Sun Q, Mitchell S, Björkman T, Hua J. 2018. Genotyping-by-sequencing of Brassica oleracea vegetables reveals unique phylogenetic patterns, population structure and domestication footprints. Hortic Res 5:38.

Swarup V, Brahmi P. 2005. Cole crops. Plant Genetic Resources: Horticultural Crops:75–88.

(the Elder.) P, Rackham H. 1950. Natural History with an English Translation: Vol 5: Libri XVII-XIX. Harvard University Press

Tomlinson P, Hall A. 1996. A review of the archaeological evidence for food plants from the British Isles: an example of the use of the Archaeobotanical Computer Database (ABCD). Internet Archaeology 1.

Totelin LMV. 2009. Hippocratic Recipes: Oral and Written Transmission of Pharmacological Knowledge in Fifth-And Fourth-Century Greece. BRILL

Turner-Hissong SD, Mabry ME, Beissinger TM, Ross-Ibarra J, Chris Pires J. 2020. Evolutionary insights into plant breeding. Current Opinion in Plant Biology [Internet] 54:93–100. Available from: http://dx.doi.org/10.1016/j.pbi.2020.03.003

Tutin TG. 1964. Flora Europaea: Lycopodiaceae to Platanaceae. University Press

Van der Auwera GA, Carneiro MO, Hartl C, Poplin R, Del Angel G, Levy-Moonshine A, Jordan T, Shakir K, Roazen D, Thibault J, et al. 2013. From FastQ data to high-confidence variant calls: the genome analysis toolkit best practices pipeline. Curr. Protoc. Bioinformatics 43:11–10.

Van der Veen M. 2011. Consumption, Trade and Innovation. Africa Magna Verlag

Vavilov. 1926. On the Origin of Cultivated Plants. Food History: Critical and Primary Sources [Internet]. Available from: http://dx.doi.org/10.5040/9781474220095.ch-004

Vavilov. 1951. The Origin, Variation, Immunity and Breeding of Cultivated Plants. Soil Science [Internet] 72:482. Available from: http://dx.doi.org/10.1097/00010694-195112000-00018

Wang H, Vieira FG, Crawford JE, Chu C, Nielsen R. 2017. Asian wild rice is a hybrid swarm with extensive gene flow and feralization from domesticated rice. Genome Res. 27:1029–1038.

West ML. 2011. Studies in Greek Elegy and Iambus. Walter de Gruyter

Widén B, Andersson S, Rao G-Y, Widén M. 2002. Population divergence of genetic (co)variance matrices in a subdivided plant species, Brassica cretica: G matrix variation inBrassica cretica. J. Evol. Biol. 15:961–970.

Yang S, Zheng Z, Mao L, Li J, Chen B. 2018. Pollen morphology of selected crop plants from southern China and testing pollen morphological data in an archaeobotanical study. Veg. Hist. Archaeobot. 27:781–799.

Yonge CD. 1854. Athenaeus of Naucratis. The deipnosophists, or, Banquet of the learned of Athenæus. London: Henry G. Bohn.

Zhang B, Horvath S. 2005. A general framework for weighted gene co-expression network analysis. Stat. Appl. Genet. Mol. Biol. 4:Article17.

Zhang P. 2009. Studies on the origin of Brassica alboglabra Bailey. China Vegetables:62–65

