## Supplemental Tables for "The Evolutionary History of Wild, Domesticated, and Feral *Brassica oleracea* (Brassicaceae)"

**Table S1.** *Brassica oleracea* crop types with common name, species name, Kew cultivar group, and other used names. Illustrations by Andi Kur.

| Common Name | Species Name | Kew Cultivar Group | Other Used Names | Phenotype |
|---|---|---|---|---|
| Broccoli | *Brassica oleracea var.italica* | Botrytis Group | | 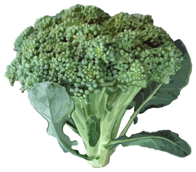 |
| Brussels spouts | *Brassica oleracea var. gemmifera* | Gemmifera Group | | 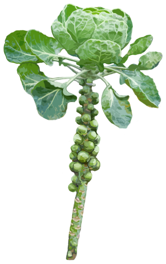 |
| Cauliflower | *Brassica oleracea var. botrytis* | Botrytis Group | | 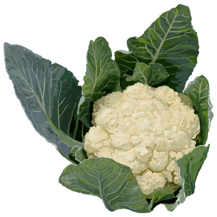 |
| Chinese White Kale | *Brassica oleracea var. alboglabra* | Alboglabra Group | Chinese broccoli, gai laan, kai lan | 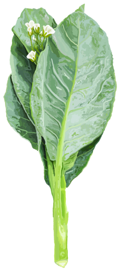 |
| Collard Greens | *Brassica oleracea var. viridis* | Acephala Group | | 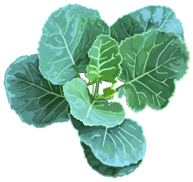 |
| Curly Kale | *Brassica oleracea var. sabellica* | Acephala Group | *Brassica oleracea* var. *acephala* | 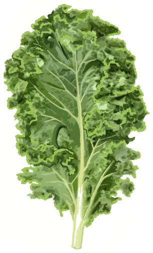 |

| | | | | |
|---|---|---|---|---|
| Giant Jersey Kale | *Brassica oleracea var. longata* | Acephala Group | Jersey Cabbage, Cow cabbage, Walking stick kale, Couve Galega, Portuguese tree kale | 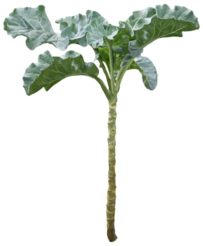 |
| Kohlrabi | *Brassica oleracea var. gongylodes* | Gongylodes Group | | 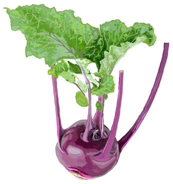 |
| Lacinato kale | *Brassica oleracea var. palmifolia* | Acephala Group | | 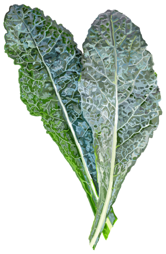 |
| Marrow Cabbage | *Brassica oleracea var. medullosa* | Acephala Group | *Brassica oleracea* var. *acephala* | 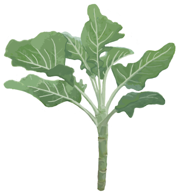 |
| Ornamental cabbage | *Brassica oleracea var. acephala* | Acephala Group | | 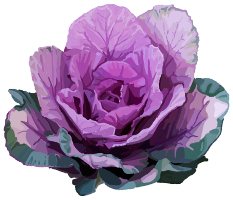 |
| Perpetual Kale | *Brassica oleracea var. ramosa* | Acephala Group | *Brassica oleracea* var. *acephala* | 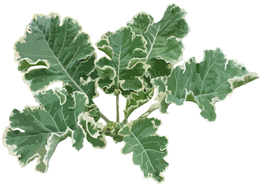 |
| Red Cabbage | *Brassica oleracea var. capitata f. rubra* | Capitata Group | | |
| Romanesco | *Brassica oleracea var. botrytis* | Botrytis Group | | 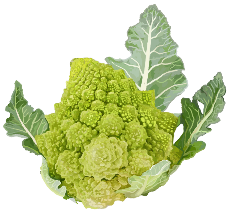 |

| | | | | |
|---|---|---|---|---|
| Savoy Cabbage | *Brassica oleracea var. capitata f. sabauda* | Capitata Group | | 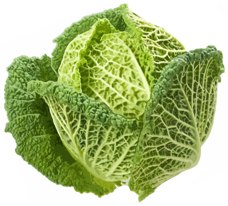 |
| Tronchuda Kale | *Brassica oleracea var. costata* | Tronchuda Group | Portuguese cabbage, seakale cabbage | 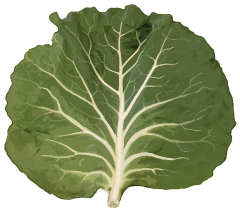 |
| White Cabbage | *Brassica oleracea var. capitata f. alba* | Capitata Group | | 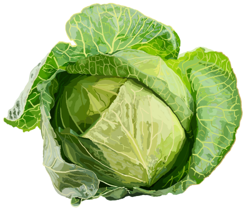 |

**Table S2.** Sample information with species, variety, cultivar, accession/collection, sample #, and SRA #. Asterisks (*) next to sample # indicate those samples that were recovered as monophyletic and used in species tree reconstruction.

| Genus | Species | Variety | Cultivar | Accession | Sample # |
|---|---|---|---|---|---|
| ***Brassica*** | *oleracea* | *virdis* | Collards | PI662796 | 180 |
| ***Brassica*** | *oleracea* | *virdis* | Collards | PI662798 | 190* |
| ***Brassica*** | *oleracea* | *virdis* | Collards | PI662804 | 184 |
| ***Brassica*** | *oleracea* | *virdis* | Collards | PI662805 | 189 |
| ***Brassica*** | *oleracea* | *virdis* | Collards | PI662806 | 181 |
| ***Brassica*** | *oleracea* | *virdis* | Collards | PI662807 | 186 |
| ***Brassica*** | *oleracea* | *virdis* | Collards | PI6628080 | 191 |
| ***Brassica*** | *oleracea* | *virdis* | Collards | PI662809 | 183* |
| ***Brassica*** | *oleracea* | *virdis* | Collards | PI662814 | 182* |
| ***Brassica*** | *oleracea* | *virdis* | Collards | PI662815 | 178* |
| ***Brassica*** | *oleracea* | *virdis* | Collards | PI6628160 | 193* |
| ***Brassica*** | *oleracea* | *virdis* | Collards | PI662817 | 179 |
| ***Brassica*** | *oleracea* | *virdis* | Collards | PI662828 | 192* |
| ***Brassica*** | *oleracea* | *virdis* | Collards | PI662831 | 185* |
| ***Brassica*** | *oleracea* | *virdis* | Collards | PI662833 | 187* |
| ***Brassica*** | *oleracea* | *virdis* | Collards | PI6628410 | 194* |
| ***Brassica*** | *oleracea* | *sabellica* | Curly Kale | IPK:BRA1491 | 54* |
| ***Brassica*** | *oleracea* | *sabellica* | Curly Kale | IPK:BRA1841 | 53* |
| ***Brassica*** | *oleracea* | *sabellica* | Curly Kale | IPK:BRA1899 | 55* |
| ***Brassica*** | *oleracea* | *sabellica* | Curly Kale | IPK:BRA2116 | 59* |
| ***Brassica*** | *oleracea* | *sabellica* | Curly Kale | IPK:BRA2187 | 50* |
| ***Brassica*** | *oleracea* | *sabellica* | Curly Kale | IPK:BRA2239 | 51* |
| ***Brassica*** | *oleracea* | *sabellica* | Curly Kale | IPK:BRA2286 | 52* |
| ***Brassica*** | *oleracea* | *sabellica* | Curly Kale | IPK:BRA2449 | 49* |
| ***Brassica*** | *oleracea* | *sabellica* | Curly Kale | IPK:BRA2456 | 48* |
| ***Brassica*** | *oleracea* | *sabellica* | Curly Kale | IPK:BRA960 | 56* |
| ***Brassica*** | *oleracea* | *sabellica* | Curly Kale | IPK:BRA963 | 57* |
| ***Brassica*** | *oleracea* | *sabellica* | Curly Kale | IPK:BRA964 | 58* |
| ***Brassica*** | *oleracea* | *sabauda* | Savoy Cabbage | Pires Collection 1 | 102 |
| ***Brassica*** | *oleracea* | *sabauda* | Savoy Cabbage | PI246117 | 100 |
| ***Brassica*** | *oleracea* | *sabauda* | Savoy Cabbage | PI343603 | 99 |
| ***Brassica*** | *oleracea* | *sabauda* | Savoy Cabbage | PI507856 | 101 |
| ***Brassica*** | *oleracea* | *ramosa* | Perpetual Kale | G30718 | 115 |
| ***Brassica*** | *oleracea* | *ramosa* | Perpetual Kale | G30724 | 116 |
| ***Brassica*** | *oleracea* | *ramosa* | Perpetual Kale | IPK:BRA1213 | 103 |
| ***Brassica*** | *oleracea* | *ramosa* | Perpetual Kale | IPK:BRA1259 | 109 |
| ***Brassica*** | *oleracea* | *ramosa* | Perpetual Kale | IPK:BRA1891 | 105 |
| ***Brassica*** | *oleracea* | *ramosa* | Perpetual Kale | IPK:BRA2382 | 112 |
| ***Brassica*** | *oleracea* | *ramosa* | Perpetual Kale | IPK:BRA2782 | 108 |
| ***Brassica*** | *oleracea* | *ramosa* | Perpetual Kale | IPK:BRA2784 | 113 |
| ***Brassica*** | *oleracea* | *ramosa* | Perpetual Kale | IPK:BRA2905 | 111 |
| ***Brassica*** | *oleracea* | *ramosa* | Perpetual Kale | IPK:BRA2952 | 104 |
| ***Brassica*** | *oleracea* | *ramosa* | Perpetual Kale | IPK:BRA2956 | 114 |
| ***Brassica*** | *oleracea* | *ramosa* | Perpetual Kale | IPK:BRA3138 | 106 |
| ***Brassica*** | *oleracea* | *ramosa* | Perpetual Kale | IPK:BRA67 | 110 |
| ***Brassica*** | *oleracea* | *ramosa* | Perpetual Kale | IPK:K9829 | 107 |
| ***Brassica*** | *oleracea* | *palmifolia* | Lacinato kale | Arias Collection 1 | 1* |
| ***Brassica*** | *oleracea* | *palmifolia* | Lacinato kale | Pires Collection 2 | 2* |
| ***Brassica*** | *oleracea* | *palmifolia* | Lacinato kale | IPK:BRA1905 | 5* |
| ***Brassica*** | *oleracea* | *palmifolia* | Lacinato kale | IPK:BRA1906 | 4* |
| ***Brassica*** | *oleracea* | *palmifolia* | Lacinato kale | IPK:BRA2846 | 6* |
| ***Brassica*** | *oleracea* | *palmifolia* | Lacinato kale | IPK:BRA3073 | 7* |
| ***Brassica*** | *oleracea* | *palmifolia* | Lacinato kale | Pires Collection 3 | 3* |
| ***Brassica*** | *oleracea* | *palmifolia* | Giant Jersey Kale | G30723 | 174 |
| ***Brassica*** | *oleracea* | *oleracea* | Wild oleracea | G30186 | 175 |
| ***Brassica*** | *oleracea* | *oleracea* | Wild oleracea | NGB16241 | 177 |
| ***Brassica*** | *oleracea* | *oleracea* | Wild oleracea | NGB21675 | 176 |
| ***Brassica*** | *oleracea* | *medullosa* | Marrow Cabbage | G30717 | 35 |
| ***Brassica*** | *oleracea* | *medullosa* | Marrow Cabbage | G30859 | 36* |
| ***Brassica*** | *oleracea* | *medullosa* | Marrow Cabbage | G30862 | 37 |
| ***Brassica*** | *oleracea* | *medullosa* | Marrow Cabbage | IPK:CR1235 | 44 |
| ***Brassica*** | *oleracea* | *medullosa* | Marrow Cabbage | IPK:CR1238 | 47 |
| ***Brassica*** | *oleracea* | *medullosa* | Marrow Cabbage | IPK:CR1256 | 43* |
| ***Brassica*** | *oleracea* | *medullosa* | Marrow Cabbage | IPK:CR1295 | 46* |

| | | | | | |
|---|---|---|---|---|---|
| ***Brassica*** | *oleracea* | *medullosa* | Marrow Cabbage | IPK:CR1317 | 42 |
| ***Brassica*** | *oleracea* | *medullosa* | Marrow Cabbage | IPK:CR1410 | 45* |
| ***Brassica*** | *oleracea* | *medullosa* | Marrow Cabbage | IPK:CR1431 | 40* |
| ***Brassica*** | *oleracea* | *medullosa* | Marrow Cabbage | IPK:CR1454 | 41* |
| ***Brassica*** | *oleracea* | *medullosa* | Marrow Cabbage | IPK:CR2545 | 39 |
| ***Brassica*** | *oleracea* | *medullosa* | Marrow Cabbage | PI662703 | 38* |
| ***Brassica*** | *oleracea* | *longata* | walking stick kale | Pires Collection 4 | 172 |
| ***Brassica*** | *oleracea* | *longata* | walking stick kale | Pires Collection 5 | 173 |
| ***Brassica*** | *oleracea* | *longata* | walking stick kale | Pires Collection 6 | 171* |
| ***Brassica*** | *oleracea* | *italica* | Broccoli | Early-Big | 168* |
| ***Brassica*** | *oleracea* | *italica* | Broccoli | Early-Big | 76* |
| ***Brassica*** | *oleracea* | *italica* | Broccoli | G21111 | 72* |
| ***Brassica*** | *oleracea* | *italica* | Broccoli | G28865 | 71* |
| ***Brassica*** | *oleracea* | *italica* | Broccoli | G30416 | 70* |
| ***Brassica*** | *oleracea* | *italica* | Broccoli | G30778 | 69* |
| ***Brassica*** | *oleracea* | *italica* | Broccoli | G30779 | 73* |
| ***Brassica*** | *oleracea* | *italica* | Broccoli | GD33-DH | 170* |
| ***Brassica*** | *oleracea* | *italica* | Broccoli | GD33-DH | 75* |
| ***Brassica*** | *oleracea* | *italica* | Broccoli | Mar34-DH | 169* |
| ***Brassica*** | *oleracea* | *italica* | Broccoli | Mar34-DH | 74* |
| ***Brassica*** | *oleracea* | *italica* | Broccoli | PI231210 | 60 |
| ***Brassica*** | *oleracea* | *italica* | Broccoli | PI662524 | 63 |
| ***Brassica*** | *oleracea* | *italica* | Broccoli | PI662525 | 68 |
| ***Brassica*** | *oleracea* | *italica* | Broccoli | PI662529 | 65 |
| ***Brassica*** | *oleracea* | *italica* | Broccoli | PI662596 | 62* |
| ***Brassica*** | *oleracea* | *italica* | Broccoli | PI662597 | 61* |
| ***Brassica*** | *oleracea* | *italica* | Broccoli | PI662674 | 64* |
| ***Brassica*** | *oleracea* | *italica* | Broccoli | PI662712 | 66* |
| ***Brassica*** | *oleracea* | *italica* | Broccoli | PI662786 | 67* |
| ***Brassica*** | *oleracea* | *gongylodes* | Kohlrabi | G30762 | 90* |
| ***Brassica*** | *oleracea* | *gongylodes* | Kohlrabi | G30935 | 84* |
| ***Brassica*** | *oleracea* | *gongylodes* | Kohlrabi | HRIGRU005389 | 167* |
| ***Brassica*** | *oleracea* | *gongylodes* | Kohlrabi | IPK:BRA1889 | 85* |
| ***Brassica*** | *oleracea* | *gongylodes* | Kohlrabi | IPK:BRA2349 | 89* |
| ***Brassica*** | *oleracea* | *gongylodes* | Kohlrabi | IPK:BRA42 | 88* |
| ***Brassica*** | *oleracea* | *gongylodes* | Kohlrabi | PI096969 | 86* |
| ***Brassica*** | *oleracea* | *gongylodes* | Kohlrabi | PI188610 | 87* |
| ***Brassica*** | *oleracea* | *gongylodes* | Kohlrabi | PI662560 | 80* |
| ***Brassica*** | *oleracea* | *gongylodes* | Kohlrabi | PI662563 | 77* |
| ***Brassica*** | *oleracea* | *gongylodes* | Kohlrabi | PI662623 | 79* |
| ***Brassica*** | *oleracea* | *gongylodes* | Kohlrabi | PI662624 | 82* |
| ***Brassica*** | *oleracea* | *gongylodes* | Kohlrabi | PI662625 | 83* |
| ***Brassica*** | *oleracea* | *gongylodes* | Kohlrabi | PI662671 | 81* |
| ***Brassica*** | *oleracea* | *gongylodes* | Kohlrabi | PI662704 | 78* |
| ***Brassica*** | *oleracea* | *gemmifera* | Brussels Sprouts | G30601 | 9* |
| ***Brassica*** | *oleracea* | *gemmifera* | Brussels Sprouts | G30867 | 12* |
| ***Brassica*** | *oleracea* | *gemmifera* | Brussels Sprouts | G30869 | 24* |
| ***Brassica*** | *oleracea* | *gemmifera* | Brussels Sprouts | G30870 | 11* |
| ***Brassica*** | *oleracea* | *gemmifera* | Brussels Sprouts | G30872 | 10* |
| ***Brassica*** | *oleracea* | *gemmifera* | Brussels Sprouts | PI209942 | 14* |
| ***Brassica*** | *oleracea* | *gemmifera* | Brussels Sprouts | PI243050 | 15* |
| ***Brassica*** | *oleracea* | *gemmifera* | Brussels Sprouts | PI244839 | 16* |
| ***Brassica*** | *oleracea* | *gemmifera* | Brussels Sprouts | PI249534 | 19* |
| ***Brassica*** | *oleracea* | *gemmifera* | Brussels Sprouts | PI261775 | 21* |
| ***Brassica*** | *oleracea* | *gemmifera* | Brussels Sprouts | PI312902 | 17* |
| ***Brassica*** | *oleracea* | *gemmifera* | Brussels Sprouts | PI343669 | 22* |
| ***Brassica*** | *oleracea* | *gemmifera* | Brussels Sprouts | PI343670 | 18* |
| ***Brassica*** | *oleracea* | *gemmifera* | Brussels Sprouts | PI343671 | 20* |
| ***Brassica*** | *oleracea* | *gemmifera* | Brussels Sprouts | PI343673 | 23* |
| ***Brassica*** | *oleracea* | *gemmifera* | Brussels Sprouts | PI365170 | 13* |
| ***Brassica*** | *oleracea* | *gemmifera* | Brussels Sprouts | PI385957 | 8* |
| ***Brassica*** | *oleracea* | *costata* | Tronchuda Kale | PI662564 | 28* |
| ***Brassica*** | *oleracea* | *costata* | Tronchuda Kale | PI662565 | 27* |
| ***Brassica*** | *oleracea* | *costata* | Tronchuda Kale | PI662566 | 31* |
| ***Brassica*** | *oleracea* | *costata* | Tronchuda Kale | PI662567 | 32 |
| ***Brassica*** | *oleracea* | *costata* | Tronchuda Kale | PI662568 | 33* |
| ***Brassica*** | *oleracea* | *costata* | Tronchuda Kale | PI662569 | 34* |
| ***Brassica*** | *oleracea* | *costata* | Tronchuda Kale | PI662659 | 30* |
| ***Brassica*** | *oleracea* | *costata* | Tronchuda Kale | PI662668 | 165* |
| ***Brassica*** | *oleracea* | *costata* | Tronchuda Kale | PI662668 | 29* |

| | | | | | |
|---|---|---|---|---|---|
| ***Brassica*** | *oleracea* | *costata* | Tronchuda Kale | PI662702 | 25* |
| ***Brassica*** | *oleracea* | *costata* | Tronchuda Kale | Pires Collection 7 | 26* |
| ***Brassica*** | *oleracea* | *capitata* | Red Cabbage | PI187232 | 159* |
| ***Brassica*** | *oleracea* | *capitata* | Red Cabbage | PI245012 | 164* |
| ***Brassica*** | *oleracea* | *capitata* | Red Cabbage | PI246046 | 163* |
| ***Brassica*** | *oleracea* | *capitata* | Red Cabbage | PI246059 | 162* |
| ***Brassica*** | *oleracea* | *capitata* | Red Cabbage | PI246060 | 161* |
| ***Brassica*** | *oleracea* | *capitata* | Red Cabbage | PI329197 | 160 |
| ***Brassica*** | *oleracea* | *capitata* | Ornamental cabbage | PI662701 | 144 |
| ***Brassica*** | *oleracea* | *capitata* | Cabbage | PI171529 | 97 |
| ***Brassica*** | *oleracea* | *capitata* | Cabbage | PI182149 | 96 |
| ***Brassica*** | *oleracea* | *capitata* | Cabbage | PI225859 | 98* |
| ***Brassica*** | *oleracea* | *capitata* | Cabbage | PI263061 | 94* |
| ***Brassica*** | *oleracea* | *capitata* | Cabbage | PI320913 | 91 |
| ***Brassica*** | *oleracea* | *capitata* | Cabbage | PI324236 | 92* |
| ***Brassica*** | *oleracea* | *capitata* | Cabbage | PI343514 | 93* |
| ***Brassica*** | *oleracea* | *capitata* | Cabbage | PI518837 | 95* |
| ***Brassica*** | *oleracea* | *botrytis* | Romanesco | Pires Collection 8 | 153* |
| ***Brassica*** | *oleracea* | *botrytis* | Romanesco | Pires Collection 9 | 155* |
| ***Brassica*** | *oleracea* | *botrytis* | Romanesco | Pires Collection 10 | 156* |
| ***Brassica*** | *oleracea* | *botrytis* | Romanesco | Pires Collection 11 | 154* |
| ***Brassica*** | *oleracea* | *botrytis* | Cauliflower | G28888 | 122* |
| ***Brassica*** | *oleracea* | *botrytis* | Cauliflower | G30435 | 126* |
| ***Brassica*** | *oleracea* | *botrytis* | Cauliflower | G30436 | 125* |
| ***Brassica*** | *oleracea* | *botrytis* | Cauliflower | G30438 | 124* |
| ***Brassica*** | *oleracea* | *botrytis* | Cauliflower | G30769 | 123* |
| ***Brassica*** | *oleracea* | *botrytis* | Cauliflower | G30857 | 121* |
| ***Brassica*** | *oleracea* | *botrytis* | Cauliflower | HRIGRU007826 | 128* |
| ***Brassica*** | *oleracea* | *botrytis* | Cauliflower | HRIGRU007826 | 152* |
| ***Brassica*** | *oleracea* | *botrytis* | Cauliflower | PI231208 | 119* |
| ***Brassica*** | *oleracea* | *botrytis* | Cauliflower | PI267724 | 127 |
| ***Brassica*** | *oleracea* | *botrytis* | Cauliflower | PI267724 | 157* |
| ***Brassica*** | *oleracea* | *botrytis* | Cauliflower | PI269312 | 117* |
| ***Brassica*** | *oleracea* | *botrytis* | Cauliflower | PI271445 | 120* |
| ***Brassica*** | *oleracea* | *botrytis* | Cauliflower | PI291996 | 118* |
| ***Brassica*** | *oleracea* | *alboglabra* | Chinese White Kale | A12 | 131* |
| ***Brassica*** | *oleracea* | *alboglabra* | Chinese White Kale | A12 | 148* |
| ***Brassica*** | *oleracea* | *alboglabra* | Chinese White Kale | Pires Collection 12 | 143* |
| ***Brassica*** | *oleracea* | *alboglabra* | Chinese White Kale | HRIGRU007543 | 130* |
| ***Brassica*** | *oleracea* | *alboglabra* | Chinese White Kale | HRIGRU007543 | 146* |
| ***Brassica*** | *oleracea* | *alboglabra* | Chinese White Kale | HRIGRU013023 | 132 |
| ***Brassica*** | *oleracea* | *alboglabra* | Chinese White Kale | HRIGRU013023 | 147 |
| ***Brassica*** | *oleracea* | *alboglabra* | Chinese White Kale | IPK:BRA1006 | 134 |
| ***Brassica*** | *oleracea* | *alboglabra* | Chinese White Kale | IPK:BRA1143 | 139 |
| ***Brassica*** | *oleracea* | *alboglabra* | Chinese White Kale | IPK:BRA1210 | 133* |
| ***Brassica*** | *oleracea* | *alboglabra* | Chinese White Kale | IPK:BRA1211 | 141* |
| ***Brassica*** | *oleracea* | *alboglabra* | Chinese White Kale | IPK:BRA1244 | 142* |
| ***Brassica*** | *oleracea* | *alboglabra* | Chinese White Kale | IPK:BRA1658 | 138* |
| ***Brassica*** | *oleracea* | *alboglabra* | Chinese White Kale | IPK:BRA169 | 137* |
| ***Brassica*** | *oleracea* | *alboglabra* | Chinese White Kale | IPK:BRA1747 | 140* |
| ***Brassica*** | *oleracea* | *alboglabra* | Chinese White Kale | IPK:BRA1909 | 136* |
| ***Brassica*** | *oleracea* | *alboglabra* | Chinese White Kale | IPK:BRA990 | 135* |
| ***Brassica*** | *oleracea* | *alboglabra* | Chinese White Kale | PI249556 | 149* |
| ***Brassica*** | *oleracea* | *alboglabra* | Chinese White Kale | PI435900 | 150* |
| ***Brassica*** | *oleracea* | *alboglabra* | Chinese White Kale | PI662520 | 151* |
| ***Brassica*** | *oleracea* | *alboglabra* | Chinese White Kale | TO1000 | 129* |
| ***Brassica*** | *oleracea* | *alboglabra* | Chinese White Kale | TO1000 | 145* |
| ***Brassica*** | *cretica* | - | - | BRA3053 | 198* |
| ***Brassica*** | *cretica* | - | - | BRA3092 | 199 |
| ***Brassica*** | *cretica* | - | - | PI662588 | 195 |
| ***Brassica*** | *cretica* | - | - | UPM6346 | 196 |
| ***Brassica*** | *hilarionis* | - | - | HRIGRU011463 | 203* |
| ***Brassica*** | *incana* | - | - | BRA1262 | 204* |
| ***Brassica*** | *incana* | - | - | BRA2918 | 205 |
| ***Brassica*** | *incana* | - | - | PI662584 | 208 |
| ***Brassica*** | *incana* | - | - | PI662591 | 209 |
| ***Brassica*** | *incana* | - | - | UPM5974 | 207* |
| ***Brassica*** | *insularis* | - | - | BRA2996 | 213* |
| ***Brassica*** | *insularis* | - | - | BRA3050 | 214* |
| ***Brassica*** | *insularis* | - | - | BRA3051 | 215* |

| | | | | | |
|---|---|---|---|---|---|
| ***Brassica*** | *insularis* | - | - | BRA3052 | 216* |
| ***Brassica*** | *insularis* | - | - | PI662587 | 212* |
| ***Brassica*** | *macrocarpa* | - | - | BRA2854 | 218* |
| ***Brassica*** | *macrocarpa* | - | - | BRA2944 | 219* |
| ***Brassica*** | *macrocarpa* | - | - | CO7031 | 220* |
| ***Brassica*** | *macrocarpa* | - | - | PI662585 | 221* |
| ***Brassica*** | *montana* | - | - | BRA1644 | 222 |
| ***Brassica*** | *montana* | - | - | GC3607-75 | 224* |
| ***Brassica*** | *rupestris* | - | - | BRA2851 | 226* |
| ***Brassica*** | *rupestris* | - | - | BRA2945 | 227* |
| ***Brassica*** | *rupestris* | - | - | BRA2992 | 228* |
| ***Brassica*** | *rupestris* | - | - | GC3822-75 | 229* |
| ***Brassica*** | *rupestris* | - | - | K10259 | 230 |
| ***Brassica*** | *rupestris* | - | - | K10260 | 231* |
| ***Brassica*** | *rupestris* | - | - | UPM6575 | 232* |
| ***Brassica*** | *villosa* | - | - | BRA1896 | 233 |
| ***Brassica*** | *villosa* | - | - | BRA2853 | 234* |
| ***Brassica*** | *villosa* | - | - | K10263 | 236* |
| ***Brassica*** | *villosa* | - | - | K10264 | 237 |
| ***Brassica*** | *villosa* | - | - | UPM6581 | 238* |

**Table S3.** WGCNA predicted gene modules with number of genes in each module, the number of annotated *Arabidopsis thaliana* genes using blast and synteny, and the percent of syntenic genes represented in the module. Module number with asterisks (*) represent the five modules with the largest percent of syntenic genes in the module.

| Module | Gene Number | # of *Arabidopsis* hits (Blast) | # of *Arabidopsis* hits (Synteny) | Percent of syntenic genes in module |
|---|---|---|---|---|
| **1** | 35981 | 20982 | 18178 | 50.5 |
| **2** | 2268 | 1263 | 954 | 42.1 |
| **3** | 2241 | 2015 | 1991 | 88.8 |
| **4** | 1051 | 868 | 805 | 76.6 |
| **5** | 1007 | 860 | 849 | 84.3 |
| **6** | 726 | 454 | 308 | 42.4 |
| **7*** | 717 | 629 | 648 | 90.4 |
| **8** | 675 | 541 | 586 | 86.8 |
| **9** | 597 | 243 | 170 | 28.5 |
| **10** | 558 | 336 | 227 | 40.7 |
| **11** | 464 | 427 | 397 | 85.6 |
| **12** | 424 | 131 | 102 | 24.1 |
| **13*** | 420 | 385 | 378 | 90.0 |
| **14** | 382 | 327 | 306 | 80.1 |
| **15** | 304 | 240 | 269 | 88.5 |
| **16** | 297 | 99 | 75 | 25.3 |
| **17** | 291 | 267 | 260 | 89.3 |
| **18** | 270 | 92 | 74 | 27.4 |
| **19** | 267 | 92 | 71 | 26.6 |
| **20** | 224 | 65 | 46 | 20.5 |
| **21** | 178 | 161 | 153 | 86.0 |
| **22** | 177 | 154 | 159 | 89.8 |
| **23** | 173 | 71 | 47 | 27.2 |
| **24** | 133 | 37 | 24 | 18.0 |
| **25** | 128 | 124 | 105 | 82.0 |
| **26** | 111 | 73 | 45 | 40.5 |
| **27** | 109 | 31 | 20 | 18.3 |
| **28** | 108 | 33 | 20 | 18.5 |
| **29** | 106 | 25 | 18 | 17.0 |
| **30*** | 91 | 84 | 85 | 93.4 |
| **31*** | 80 | 78 | 75 | 93.8 |
| **32** | 75 | 71 | 67 | 89.3 |
| **33** | 71 | 17 | 13 | 18.3 |
| **34*** | 70 | 66 | 63 | 90.0 |
| **35** | 69 | 17 | 17 | 24.6 |
| **36** | 61 | 27 | 12 | 19.7 |
| **37** | 57 | 15 | 12 | 21.1 |
| **38** | 54 | 24 | 14 | 25.9 |
| **39** | 49 | 13 | 12 | 24.5 |
| **40** | 47 | 20 | 11 | 23.4 |
| **41** | 45 | 14 | 9 | 20.0 |
| **42** | 44 | 14 | 8 | 18.2 |
| **43** | 44 | 7 | 8 | 18.2 |
| **44** | 44 | 12 | 9 | 20.5 |
| **45** | 40 | 39 | 33 | 82.5 |
| **46** | 39 | 11 | 7 | 17.9 |
| **47** | 37 | 14 | 9 | 24.3 |
| **48** | 34 | 8 | 7 | 20.6 |

**Table S4.** Top five WGCNA modules with largest percent of syntenic genes represented in the module with corresponding annotated GO biological process with the largest fold enrichment, and p-value.

| Module | GO Biological Process | Fold Enrichment | p-value |
|---|---|---|---|
| **7** | branched-chain amino acid catabolic process | 17.86 | 1.32E-03 |
| | phototropism | 16.58 | 1.29E-02 |
| | detection of abiotic stimulus | 14.21 | 2.79E-02 |
| | detection of external stimulus | 14.21 | 2.79E-02 |
| | fatty acid beta-oxidation | 12.55 | 1.03E-02 |
| **13** | mitochondrial transcription | 93.62 | 4.82E-02 |
| | snoRNA 3'-end processing | 53.5 | 2.37E-05 |
| | polyadenylation-dependent snoRNA 3'-end processing | 49.93 | 1.04E-02 |
| | endonucleolytic cleavage involved in rRNA processing | 48.01 | 6.80E-04 |
| | U4 snRNA 3'-end processing | 45.39 | 1.41E-02 |
| **30** | suberin biosynthetic process | > 100 | 3.52E-08 |
| | phenylpropanoid biosynthetic process | 54.01 | 5.32E-10 |
| | phenylpropanoid metabolic process | 43.61 | 3.23E-09 |
| | secondary metabolite biosynthetic process | 33.01 | 3.43E-08 |
| | pectin catabolic process | 26.94 | 4.29E-03 |
| **31** | cytidine to uridine editing | 75.81 | 1.25E-03 |
| | mitochondrial RNA modification | 60.3 | 1.14E-04 |
| | base conversion or substitution editing | 58.96 | 3.08E-03 |
| | mitochondrial mRNA modification | 56.86 | 3.51E-03 |
| | RNA modification | 42.34 | 6.27E-37 |
| **34** | wax biosynthetic process | > 100 | 3.59E-12 |
| | wax metabolic process | > 100 | 4.61E-12 |
| | fatty acid derivative biosynthetic process | > 100 | 7.43E-12 |
| | cuticle development | > 100 | 1.45E-11 |
| | very long-chain fatty acid biosynthetic process | > 100 | 2.62E-04 |

**Table S6:** Archaeological *Brassica* reports from Europe and the Eastern Mediterranean. BP = years before the present (1950).

| Date | Countries | Notes | References |
|---|---|---|---|
| **Middle Bronze Age ca. 3550 - 3350 BP** | Austria, Friaga | 3 charred seeds, hilltop settlement in the southern Alps. Unusually early date, but no reason to suspect contamination from later occupation etc. | *1* |
| **ca. 3250 - 2970 BP** | Gibala, NW Syria | C14 dated seeds from pottery vessels associated with a destruction layer. Seeds have been SEM; Seeds could be *Brassica cretica*. | *2* |
| **La Tène 2400 - 2050 BP** | Switzerland/France | Preservation not specified but is absent in Roman samples. Refers to identification problems. | *3* |
| **Late Iron Age 2350 - 2050 BP** | Germany | Review of various finds including *Brassica oleracea* from the German Late Iron Age | *4* |
| **Roman-Iron Age 1950 - 1650 BP** | Egypt, Mons Claudianus | charred/desiccated seeds resembling *Brassica oleracea* | *5* p.200 |
| **1950 – 1550 BP** | Italy, Vado Ligure, Liguria | possible seeds found in Roman well | *6* |
| **1907-1540 BP** | England | recorded from 3 military and 2 town sites | *7* |
| **Roman 1850-1650 BP** | Germany, Otterbach, Kaiserslautern | 11 waterlogged seeds of probable *Brassica* cf. *oleracea* from a well in Otterbach | *8* |
| **1550-850 BP** | Sweden, Denmark, Netherlands | seeds of *Brassica oleracea* | *9, 10* |
| **1150-700 BP** | Czech Republic, Žatec | Finds of waterlogged *Brassica* cf. *oleracea* | *11* |

| | | | |
|---|---|---|---|
| **1050-850 BP** | Great Britain, York | Finds of waterlogged *Brassica* cf. *oleracea* | *12* |
| **1050-850 BP** | Great Britain, York | Finds of waterlogged *Brassica* cf. *oleracea* | *13* |
| **850 BP** | Denmark | One to four chance finds from cultural layers | *14* |
| **Early Medieval 850-750 BP** | England, Raunds, Northants | Residues of *Brassica* leaves identified within pottery sherds. | *15, 16* |
| **727-650 BP** | Montgomery, Powys, Wales | Single waterlogged seed comparable to *Brassica oleracea* from pit fill associated with castle | *17* |
| **750 - 650 BP** | Germany, Einbeck / Greifswald | Archaeobotanical material including mid. 14th century sewers. [73] p.150 | *18, 19* |
| **Late Middle Ages 650 - 450 BP** | Denmark, København | Archaeobotanical finds | *20* |
| **Medieval 550 - 450 BP** | Hungrary, Budapest | Waterlogged seeds in well | *21* (Table 6.1) |
| **650-350 BP** | Germany, Stralsund, Kiel; Lüneburg; Greifswald | Mainly waterlogged seeds of *Brassica* cf. *oleracea* | *22, 23* |
| **450- 350 BP** | Netherlands, Haarlem | Uncertain identification of mineralised seed of *Brassica* cf. *oleracea* | *24* |
| **450- 350 BP** | Germany, Rostock | archaeobotanical finds | *25* |
| **350-250 BP** | Belgium, Antwerpen | Seeds found in contents of 17th Century waste pit at Antwerpen | *26* |

| | | | |
|---|---|---|---|
| **350-250 BP** | Hungary, Buda | Probable charred seed of cabbage (*Brassica* cf. *oleracea*) found within waste pit | *27* |

**Supplementary References**


1. Schmidl, A., Oeggl, K. Subsistence strategies of two hilltop settlements in the Eastern Alps - Friaga/Bartholomäberg (Vorarlberg, Austria) and Ganglegg/Schluderns (South Tyrol, Italy). *Veget. Hist. Archaeobot.* 14, 303–312 (2005).

2. Kaniewski, D. *et al*., The Sea Peoples, from cuneiform tablets to carbon dating**.** *PLoS ONE* **6**(6), e20232 (2011).

3. Jacomet, S., Vandorpe, (Trans. Martinoli, D.) Plantes anciennes et nouvelles. La région du Rhin supérieur et l'Allemagne du Sud-Ouest, in Reddé *et al*., Eds., Aspects de la Romanisation dans l'Est de la Gaule. (Bibracte, 2011). pp. 345-360.

4. Willerding, U. Obstarten und Nüsse; Garen-, Obst- und Weinbau; Gartenbau, Obstbau, Weinbau in Benecke, N., Donat, P., Gringmuth-Dallmer, E., Willerding, U. Eds., *Frühgeschichte der Landwirtschaft in Deutschland Beitr Ur- u Frühgesch Mitteleuropa 14. Langenweißbach*, (2003), pp.30-31, pp. 162-172, pp. 258-262.

5. van der Veen, M. The botanical evidence. in Maxfield, V.A., Peacock, D.P.S.. *Mons Claudianus. Survey and excavation 1987-1993. Vol 2. Excavations: Part 1. Fouilles* (IFAO [Inst Franç Archéol Orientale] 43. Paris, 2001), pp. 174-222; appendix 1-2; fig. 8.1-8.13.

6. Arobba, D., Bulgarelli, F., Siniscalco, C., Caramiello, R. Roman landscape and agriculture on the Ligurian coast through macro and microremains from a Vada Sabatia well (Vado Ligure, Italy). *Environmental Archaeology* **18**(2), 114-131 (2013).

7. van der Veen, M., Livarda, A., Hill, A. New food plants in Roman Britain – dispersal and social access. *Environmental Archaeology* **13**(1), 11-36 (2008).

8. Weitholdd, J. Hirse, Hanf und Hohldotter: Pflanzenfunde aus einem römischen Brunnen in Otterbach, Kr. Kaiserslautern, in Stobbe, A., Tegtmeier, U. Eds., *Verzweigungen: Eine Würdigung für A. J. Kalis und J. Meurers-Balke*. (Verlag Dr. Rudolf Habelt GmbH, Bonn, 2012).

9. Hansson, A.-M. Vikingabotanik. in Pettersson, B. *et al*., Eds., *Människan och naturen.Etnobiologi i Sverige* 1. (Stockholm, 2001).

10. Heimdahl, J. Barbariska trädgårdsmästare.Nya perspektiv på hortikulturen i Sverige fram till 1200-talets slut. *Fornvännen* **105**, 265-280 (2010).

11. Kočár, P., Čech, P., Kozáková, R., Kočárová, R. Environment and economy of the Early Medieval settlement in Žatec. *Interdiciplinaria Archaeologica, Natural Sciences in Archaeology (IANSA)* **1**(1-2), 45-60 (2010).

12. Godwin, H., Bachem, K. Appendix III Plant Materials. 109-13. in Richardson, K.M. Excavations in Hungate, York. *Archaeol. J.* **116** (for 1959), 51-114 (1961).
13. Hall, A.R., Williams, D., Greig, J.R.A. [plant remains]. 157-225 & fiche. in Hall, A.R., Kenward, H.K., Williams, D., Greig, J.R.A. Environment and living conditions at two Anglo-Scandinavian sites. The Archaeology of York AY **14**(4). London: CBA. 157-240 and fiche 1 (1983).

14. Jensen, H.-A. Catalogue of late- and post-glacial macrofossils of Spermatophyta from Denmark, Schleswig, Scania, Halland, and Blekinge dated 13,000 B.P. to 1536 A.D. *Danmarks Geologiske Undersøgelser Serie A*, nr. 6 1-95. (København, Reitzels forlag, 1985).

15. Evershed, R.P., Heron, C., Charters, S., Goad, L.J. The survival of food residues: new methods of analysis, interpretation and application. *Proc. Brit. Acad.* **77**, 189–208 (1992).

16. Evershed, R.P., Amot, K.L., Collister, J., Eglinton, G., Charters, S. Application of isotope ratio monitoring gas chromatography-mass spectrometry to the analysis of organic residues of archaeological origin. *Analyst* **119**, 909–14 (1994).

17. Greig, J., Girling, M., Skidmore, P. The plant and insect remains. 60-71 in Barker, P., Higham, R. *Hen Domen Montgomery. A timber castle on the English-Welsh Border.* (I. Roy. Archaeol. Inst., London, 1982).

18. Wiethold, J. Giff in de schottele. Strowe dar peper up …Botanische Funde als Quellen zur mittel-alterlichen Ernährungs- und Umweltgeschichte in Einbeck. in Heege A. (Hrsg.), *Einbeck im Mittelalter : eine archäologisch-historische Spurensuche*. Studien zur Einbecker Geschichte 17 (Oldenburg 2002), pp. 240-246.

19. Ansorge, J., Igel, K., Schäfer, H., Wiethold, J. Ein Holzschacht aus der Baderstr. 1 in Greifswald. Aus der materiellen Alltagskultur der sozialen Oberschicht einer Hansestadt in der 2. Hälfte des 14. Jahrhunderts. Bodendenkmalpflege in Mecklenburg-Vorpommern, Jahrbuch 50, 2002, 119-157 (2003).

20. Moltsen, A.S.A. Arkæobotaniske undersøgelser fra middelalderbyen København - metodik og udvalgte eksempler in Viklund, K. Ed., *Nordic archaeobotany - NAG 2000 in Umeå*. Archaeology and Environment 15. (Univ. Umeå, Umeå, 2002). pp. 173-180.

21. Zoltán, A., Sárgadinnye, A. (Cucumis melo) Archeogenetikája, ITS- és SSR – Heterogenitása Egy 15. századi Lelettől a Mai Fajtákig, Unpublished Doctoral Thesis. Gödöllő. (2006).

22. Wiethold, J. Archäobotanische Ergebnisse. in Fries, H., Wiethold, J. *Bemerkenswertes aus Stralsunds Altstadt die Grabung Apollonienmarkt 6 und ihre Ergebnisse*. (Archäol Ber Mecklenburg-Vorpommern 10, 2003), pp. 220-247.

23. Wiethold, J. Archäobotanische Untersuchungen zur Ernährungs- und Wirtschaftsgeschichte des Mittelalters und der frühen Neuzeit. in Noël, R., Paquay, I., Sosson, J.-P. Eds., *Au-delà de l'écrit. Les hommes et leurs vécus matériels au Moyen Âge à la lumière des sciences et des techniques. Nouvelles perspectives*. (Louvain-la-Neuve, 2003), pp. 461-499.

24. Brinkkemper, O. Plantenresten uit beerputten aangetroffen op een bouwlocatie aan het Spaarne te Haarlem. Een ‚rijke' informatiebron! *Haarlems Bodemonderzoek* **36**, 104-132 (2002).

25. Wiethold, J. Ernährung und Umwelt im spätmittelalterlichen Rostock. Archäobotaniscje Ergebnisse der Analyse zweier Kloaken in der Kröpeliner Straße 55-56, Kuhstraße. *Bodendenkmalpflege Mecklenburg-Vorpommern, Jahrbuch 1999* **47**, 351-378 (2000).

26. Cooremans, B. Het macrobotanisch onderzoek: een speurtocht naar plantaardige (voedsel) resten. in Veeckman, J., van Hoof, W., Cooremans, B., Ervynck, A., van Neer, W. Eds., *De inhoud van de afvalput van de Groote Schalien Loove: speuren naar de 17de eeuwsde bewoners.* BRABOM Berichten en rapporten over het Antwerps. Bodemonderzoek en Monumentenzorg 3. (Antwerpen, Stad Antwerpen, 2000), pp. 131-139.

27. Tóth, A.J., Daróczi-Szabó, L., Kovács, Zs.E., Gál, E., Bartosiewicz, L. In the light of the crescent moon: reconstructing environment and diet from an Ottoman Period deposit in Sixteenth to Seventeenth Century Hungary. in Peres, T., Van Derwarker, A. Eds., *Integrating Zooarchaeology and Paleoethnobotany*. (Springer, New York, 2010), pp. 245-280.

**Table S7:** Literary and artistic sources covering the Classical Greek, Roman, and medieval and post-medieval sources. BP = years before the present (1950).

| Date | Countries | Notes | References |
|---|---|---|---|
| **2500-2000 BP** | Ionian Greece | Earliest textual reference. Refers to a "seven-leaved" cabbage in iambic verse. Perhaps as an oath or offering. | Hipponax, Fr.104 [1] p. 145 |
| **2410-2320 BP** | Classical Greece | Refers to cabbage in a few medicinal recipes | Hippocrates, *Nat Mul.* 32, *Mul 1.78* [2] p. 75 |
| **ca. 2330-2250 BP** | Classical Greece | 3 varieties smooth, parsley-leaved and salty. Last has delicate taste and grows in Eretria, Cyme and Rhodes (and in Cnidos and Ephesus). | Eudemus f. 85, Quoted in Athenaeus *The Deipnosophists* 9.9 [3] pp. 582-3 |
| **ca. 2320-2238/2235 BP** | Classical Greece | 3 varieties: smooth ?seedless (capita), larger leafed, sweeter parsley/curly-leaved, and a wild type bitter taste with many branches and smaller leaves may be *B. cretica* (see [61] p. 20) | Theophrastus *Historia Plantarum* VII. iv .4-6 [4] pp. 84-85 Athenaeus The Deipnosophists 9.9 [3] pp. 582-3 |
| **ca. 2150-2250 BP** | Cyclades, Greece | States the finest cabbages grow in Cyme, but bitter in Alexandria. Seed brought from Rhodes produces a sweet cabbage than after a year taste degenerates. | Diphilus of Siphnus (the Siphnian). Athenaeus The Deipnosophists 9.9 [3] pp. 582-3 |
| **ca. 2150-2050 BP** | Greece + | Smooth leaved, sometimes found wild. Also a curly leaved and one of reddish color. | Nicander Georgics. f.85. Athenaeus The Deipnosophists 9.9 [3] pp. 582-3 |
| **ca. 2110 BP** | Classical Roman Italy | Lists smooth-leaved, curly leaved (apiaca), a small stalked, pungent tender variety and wild type | Cato, Marcus[5] De Agricultura. Capitula CLVI-CLVII:156-157, 1 pp. 144-151 |

| | | | |
|---|---|---|---|
| **1873-1871 BP** | Classical Roman Italy | List a large number of varieties. | [6] 41.19. |
| **1700-1500 BP** | Egypt | List of Monastic sources from Egyptian papyri | Oxyrhynchus Papyri XIV 1656 (BL), [7] |
| **1600-1500 BP** | Roman | Cited in nine recipes attributed to Apicius: dealing with Cimæ (cymae) & Coliculi (refers to sprouts, young cabbage and soft cabbage) | [8] Apicus III, ix.87-92, x.94, xii.99 & xv.103 |
| **1180-1150 BP** | France/ Germany | Capitularies of Charlemagne Caulos & Ravacaulos (Kohlrabi ) | Capitulare de Villis [9] p. 90 |
| **Medieval 1150-650 BP** | Poland, Krakow | archaeobotanical and literary sources. See [88] for identification criteria (also [87]) | [10, 11] |
| **Medieval Spain ca. 990-600 BP** | Spain | Translations of qannabit & kurunb would indicate cauliflower & cabbage from *c.*960 AD | [12] |
| **750-550 BP** | France/ Switzerland | Le Viandier de Taillevent: Collection of medieval recipes. Mentioned in one recipe. | [13] |
| **Late Middle Ages 650-450 BP** | Czech Republic | Reference to cabbage, garlic and onion in cooking and medicine | [14] |
| **Late Middle Ages 650-450 BP** | Estonia, Tartu | Written and artistic depictions of *Brassica oleracea* | [15, 16] |
| **450-250 BP** | Flanders and Netherlands | Depictions of cauliflower and cabbages in art. | [17] |

| | | | |
|---|---|---|---|
| **450-350 BP** | Russia | | Sil'vestr's Domostroi |
| **396 BP** | Flemish | lists 5 types of cabbage - white, red, savoy, Roosken (pale red) and curly kale | [18] |
| **370 BP** | England | Refers to time of sowing and Ionians holding them as sacred | [19] |
| **303 BP** | Denmark | *Brassica oleracea capita* referred to as garden plant in *Horticultura Danica* | [20] |

### Supplementary References


1. West, M.L. *Studies in Greek Elegy and Iambus*. (De Gruyter, 1974).

2. Totelin, L.M.V. Ed., *Hippocratic recipes: oral and written transmission of pharmacological knowledge in Fifth- and Fourth-century Greece*. (Brill., 2009).

3. Athenaeus of Naucratis *c.*3rd C AD, *The Deipnosophists: Banquet of the Learned of Athnaeus Volume II.* (Trans. Yonge). (Henry G. Bohn, London, 1854).

4. Theophrastus, *Historum Plantarum*. Enquiry into Plants, Volume II: Books 6-9: On Odours. Weather Signs. Trans. Hort, A.F. (*De Causis Plantarum*). (Loeb Classical Library 79, 1916).

5. Hooper, A.D., Ash, H.B. *Cato and Varro: On Agriculture.* Loeb Classical Library, No. 283. (Harvard Univ. Press, 1934).

6. Pliny, the Elder, AD 77-79 *Natural History*, Volume V: Books 17-19. (Trans. Rackham, H.). (Loeb Classical Library 371, 1950).

7. Harlow, M., Smith, W. Between fasting and feasting: the literary and archaeobotanical evidence for monastic diet in Late Antique Egypt, *Antiquity* **75**(290), 758–768 (2001).

8. Vehling, J.D. *Apicius. Cookery and Dining in Imperial Rome*. Trans. Joseph Dommers Vehling. (Walter M. Hill, Chicago, 1936).

9. Boretius, A. Ed., *Capitularia regum Francorum* I, MGH Legum Sectio II (Hanover, 1883), no. 32, pp. 82-91.

10. Wieserowa, A. Plant remains from the early and late Middle Ages found in the settlement layers of the main market square in Cracow. *Acta Palaeobotanica* **20**(2), 137-212 (1979).

11. Zemanek, A., Wasylikowa, K. Historia botaniki i archeobotanika w poszukiwaniu danycho użytkowaniu roślin w średniowiecznym Krakowie. *Analecta, Studia i Materiały z Dziejów Nauki*, V **1**(9). 123–138 (1996).

12. Harvey, J. Plants of Moorish Spain: a fresh look. *Garden History* **20**(1), 71-82 (1992).

13. Prescott, J. [1315-1395] *le Viandier de Taillevent: 14th Century Cookery, Based on the Vatican Library Manuscript Translated into English by James Prescott*. (Hypatia Press, Eugene, ed. 2, 1989).

14. Beranová, Zelí, cibule a česnek v kuchyni a v medicíně do konce 16. Století. *Sborník Západčeského muz Plzni [Plizen] řada History* **16**, 185-194 (2002).

15. Sillasoo, I. Gardens and garden products in Medieval Turku, Estonia, in Viklund, K. Ed., *Nordic archaeobotany* - Archaeology and Environment 15. (Univ. of Umeå, Umeå, 2002), pp. 181-192.

16. Sillasoo, I. Plant depictions in late medieval religious art, in Laszlovszky, J., Szabó, P. Eds., *People and Nature in Historical Perspective*. (Central European Univ., Budapest, 2003), pp. 377-393.

17. Zeven, A.C., Brandenburg, W.A. Use of paintings from the 16th to 19th centuries to study the history of domesticated plants. *Economic Botany* **40**(4), 397-408 (1986).

18. Dodoens, R. *Cruydeboeck*. (Antwerpen, 1554).

19. Tusser, T. (1580). *Five Hundred Pointes of Good Husbandrie*. (English Dialect Society, London, 1878).

20. Block, H.R. (1647) *Ældste danske Havebog, Horticultura Danica*. Copenhagen (Wormianum, Aarhus, 1984).

**Table S8.** Area under the receiver operating characteristic curve (AUC) values for Maxent environmental niche model runs for putative wild relatives of *Brassica oleracea* crops.

| **Species** | **AUC** |
|---|---|
| ***B. hilarionis*** | 0.988 |
| ***B. insularis*** | 0.954 |
| ***B. macrocarpa*** | 0.982 |
| ***B. montana*** | 0.972 |
| ***B. oleracea*** | 0.989 |
| ***B. rupestris*** | 0.979 |
| ***B. villosa*** | 0.977 |
| ***B. incana*** | 0.967 |
| ***B. cretica*** | 0.988 |
