## Supplemental Figures for "The Evolutionary History of Wild, Domesticated, and Feral *Brassica oleracea* (Brassicaceae)"

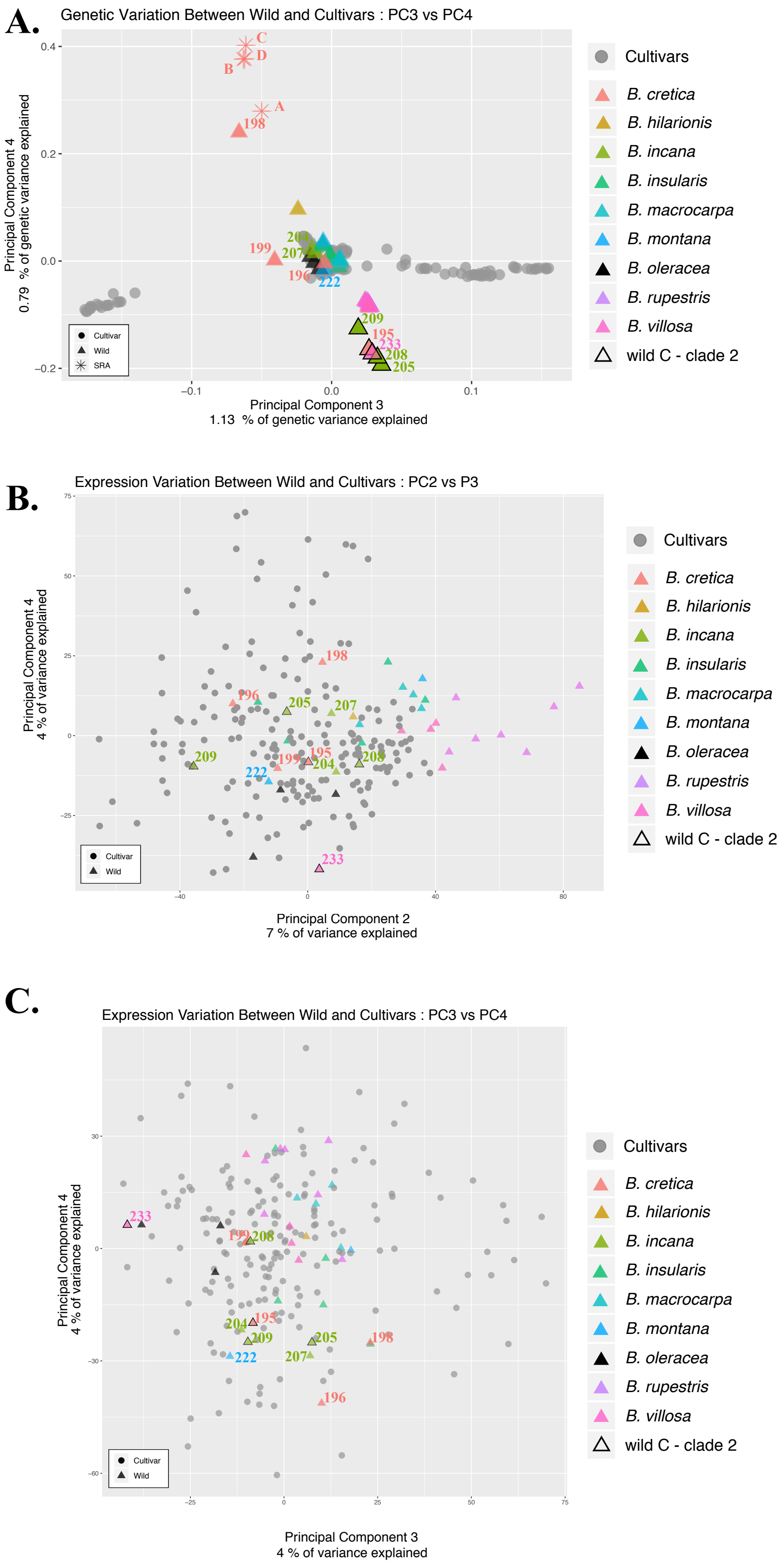
A.
Genetic Variation Between Wild and Cultivars : PC3 vs PC4
Principal Component 4
0.79 % of genetic variance explained
Principal Component 3
1.13 % of genetic variance explained
Cultivar
Wild
SRA
A
B
C
D
198
199
207
204
196
222
209
195
233
208
205
Cultivars
B. cretica
B. hilarionis
B. incana
B. insularis
B. macrocarpa
B. montana
B. oleracea
B. rupestris
B. villosa
wild C - clade 2
B.
Expression Variation Between Wild and Cultivars : PC2 vs P3
Principal Component 4
4 % of variance explained
Principal Component 2
7 % of variance explained
Cultivar
Wild
198
196
205
207
209
199
195
204
208
222
233
Cultivars
B. cretica
B. hilarionis
B. incana
B. insularis
B. macrocarpa
B. montana
B. oleracea
B. rupestris
B. villosa
wild C - clade 2
C.
Expression Variation Between Wild and Cultivars : PC3 vs PC4
Principal Component 4
4 % of variance explained
Principal Component 3
4 % of variance explained
Cultivar
Wild
233
199
208
204
195
209
222
205
207
198
196
Cultivars
B. cretica
B. hilarionis
B. incana
B. insularis
B. macrocarpa
B. montana
B. oleracea
B. rupestris
B. villosa
wild C - clade 2

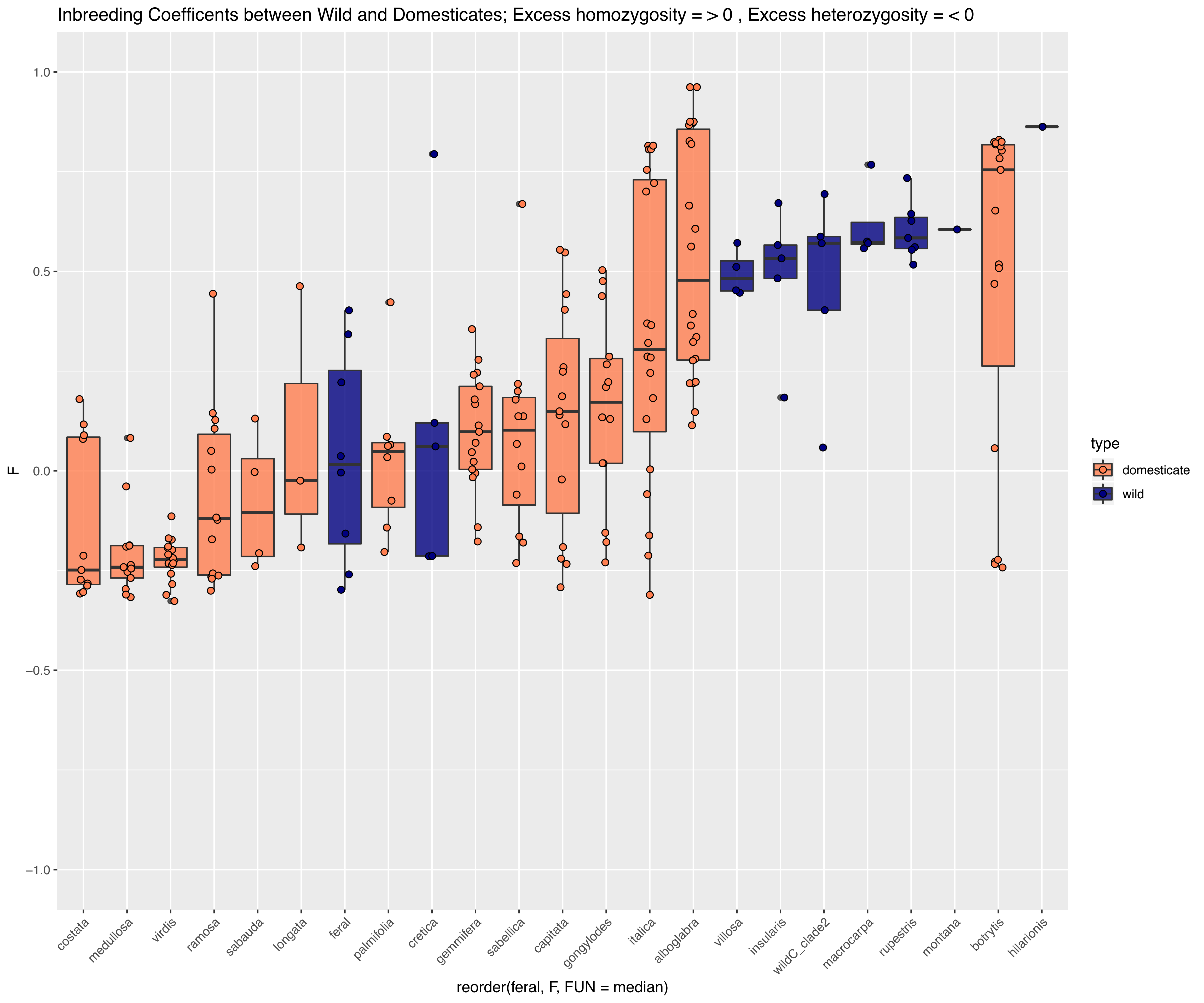

Inbreeding Coefficents between Wild and Domesticates; Excess homozygosity = > 0 , Excess heterozygosity = < 0
F
1.0
0.5
0.0
−0.5
−1.0
costata
medullosa
virdis
ramosa
sabauda
longata
feral
palmifolia
cretica
gemmifera
sabellica
capitata
gongylodes
italica
alboglabra
villosa
insularis
wildC_clade2
macrocarpa
rupestris
montana
botrytis
hilarionis
reorder(feral, F, FUN = median)
type
domesticate
wild

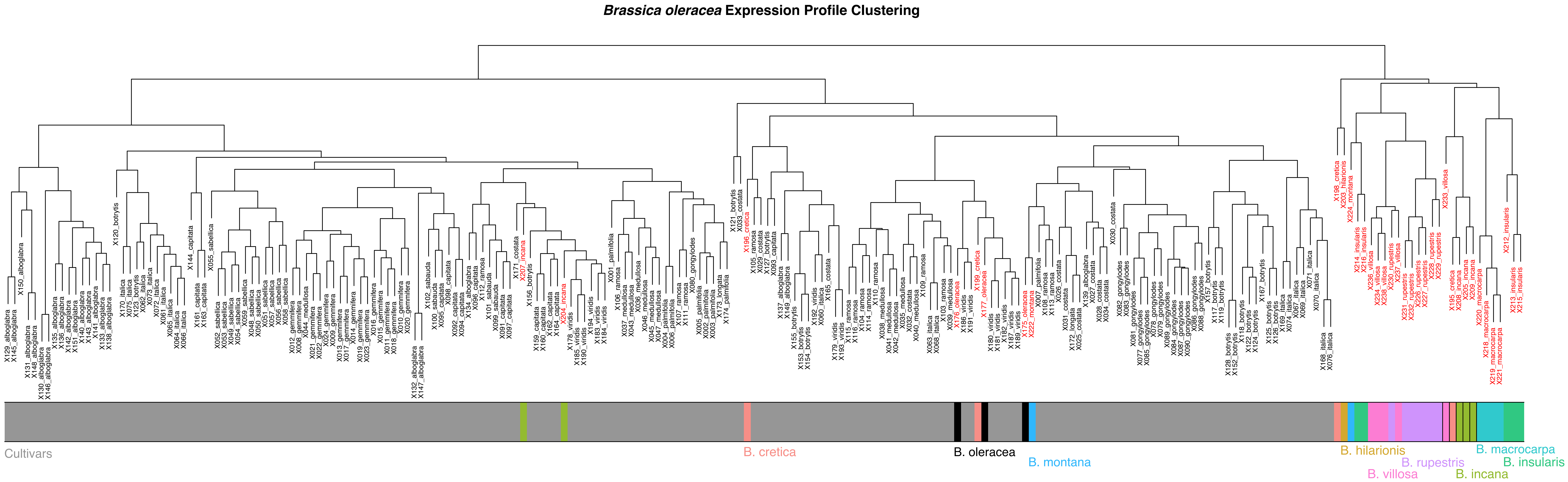
Brassica oleracea Expression Profile Clustering
Cultivars
B. cretica
B. oleracea
B. montana
B. hilarionis
B. rupestris
B. villosa
B. incana
B. macrocarpa
B. insularis

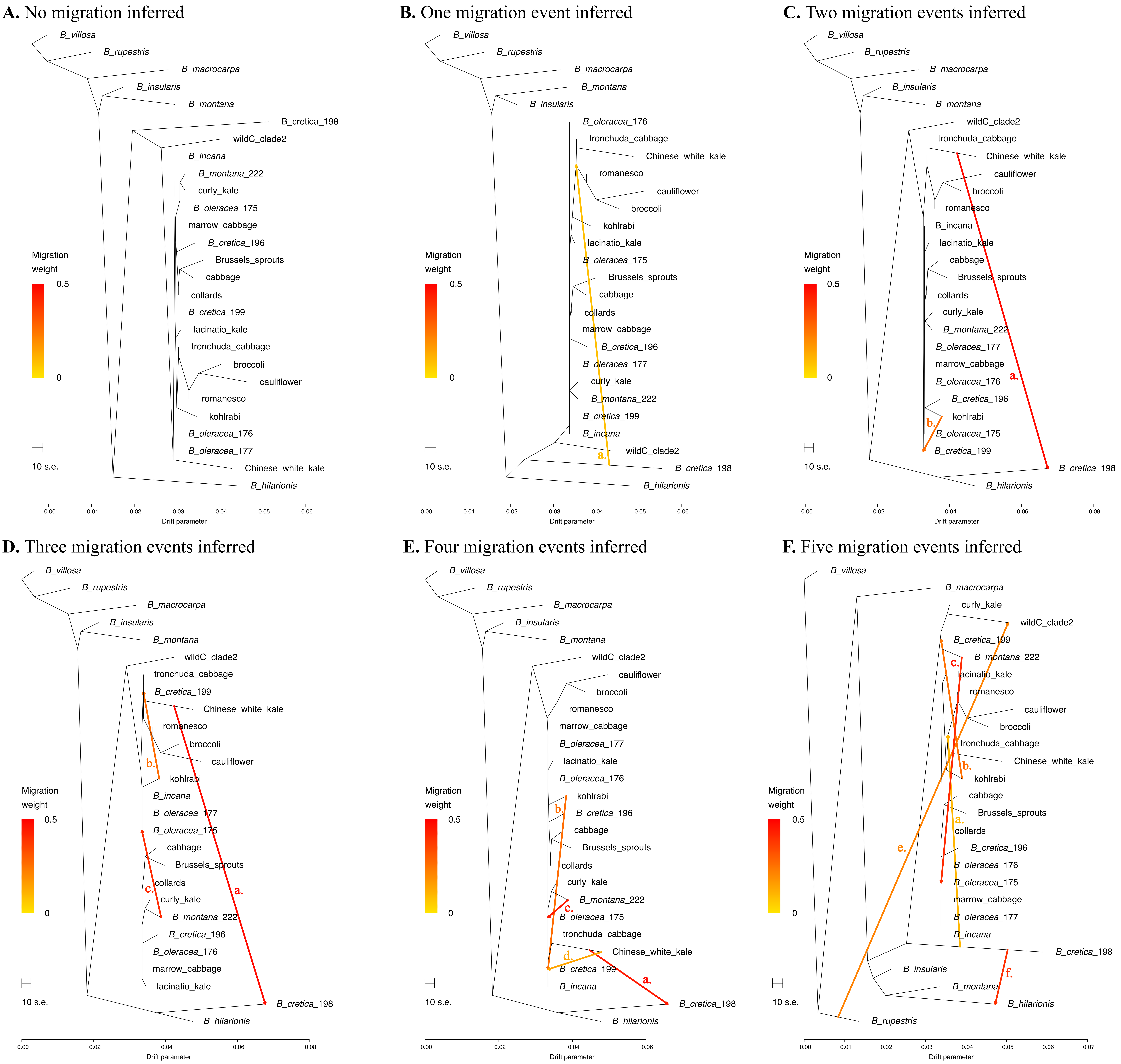
A. No migration inferred
B_villosa
B_rupestris
B_macrocarpa
B_insularis
B_montana
B_cretica_198
wildC_clade2
B_incana
B_montana_222
curly_kale
B_oleracea_175
marrow_cabbage
B_cretica_196
Brussels_sprouts
cabbage
collards
B_cretica_199
lacinatio_kale
tronchuda_cabbage
broccoli
cauliflower
romanesco
kohlrabi
B_oleracea_176
B_oleracea_177
Chinese_white_kale
B_hilarionis
Migration weight
0.5
0
10 s.e.
0.00 0.01 0.02 0.03 0.04 0.05 0.06
Drift parameter
B. One migration event inferred
B_villosa
B_rupestris
B_macrocarpa
B_montana
B_insularis
B_oleracea_176
tronchuda_cabbage
Chinese_white_kale
romanesco
cauliflower
broccoli
kohlrabi
lacinatio_kale
B_oleracea_175
Brussels_sprouts
cabbage
collards
marrow_cabbage
B_cretica_196
B_oleracea_177
curly_kale
B_montana_222
B_cretica_199
B_incana
a.
wildC_clade2
B_cretica_198
B_hilarionis
Migration weight
0.5
0
10 s.e.
0.00 0.01 0.02 0.03 0.04 0.05 0.06
Drift parameter
C. Two migration events inferred
B_villosa
B_rupestris
B_macrocarpa
B_insularis
B_montana
wildC_clade2
tronchuda_cabbage
Chinese_white_kale
cauliflower
broccoli
romanesco
B_incana
lacinatio_kale
cabbage
Brussels_sprouts
collards
curly_kale
B_montana_222
B_oleracea_177
marrow_cabbage
B_oleracea_176
a.
B_cretica_196
b.
kohlrabi
B_oleracea_175
B_cretica_199
B_cretica_198
B_hilarionis
Migration weight
0.5
0
10 s.e.
0.00 0.02 0.04 0.06 0.08
Drift parameter
D. Three migration events inferred
B_villosa
B_rupestris
B_macrocarpa
B_insularis
B_montana
wildC_clade2
tronchuda_cabbage
B_cretica_199
Chinese_white_kale
romanesco
broccoli
b.
cauliflower
kohlrabi
B_incana
B_oleracea_177
B_oleracea_175
cabbage
Brussels_sprouts
collards
c.
a.
curly_kale
B_montana_222
B_cretica_196
B_oleracea_176
marrow_cabbage
lacinatio_kale
B_cretica_198
B_hilarionis
Migration weight
0.5
0
10 s.e.
0.00 0.02 0.04 0.06 0.08
Drift parameter
E. Four migration events inferred
B_villosa
B_rupestris
B_macrocarpa
B_insularis
B_montana
wildC_clade2
cauliflower
broccoli
romanesco
marrow_cabbage
B_oleracea_177
lacinatio_kale
B_oleracea_176
kohlrabi
b.
B_cretica_196
cabbage
Brussels_sprouts
collards
curly_kale
B_montana_222
c.
B_oleracea_175
tronchuda_cabbage
d.
Chinese_white_kale
B_cretica_199
a.
B_incana
B_cretica_198
B_hilarionis
Migration weight
0.5
0
10 s.e.
0.00 0.02 0.04 0.06
Drift parameter
F. Five migration events inferred
B_villosa
B_macrocarpa
curly_kale
wildC_clade2
B_cretica_199
c.
B_montana_222
lacinatio_kale
romanesco
cauliflower
broccoli
tronchuda_cabbage
Chinese_white_kale
b.
kohlrabi
cabbage
Brussels_sprouts
a.
collards
e.
B_cretica_196
B_oleracea_176
B_oleracea_175
marrow_cabbage
B_oleracea_177
B_incana
B_cretica_198
B_insularis
f.
B_montana
B_hilarionis
B_rupestris
Migration weight
0.5
0
10 s.e.
0.00 0.01 0.02 0.03 0.04 0.05 0.06 0.07
Drift parameter

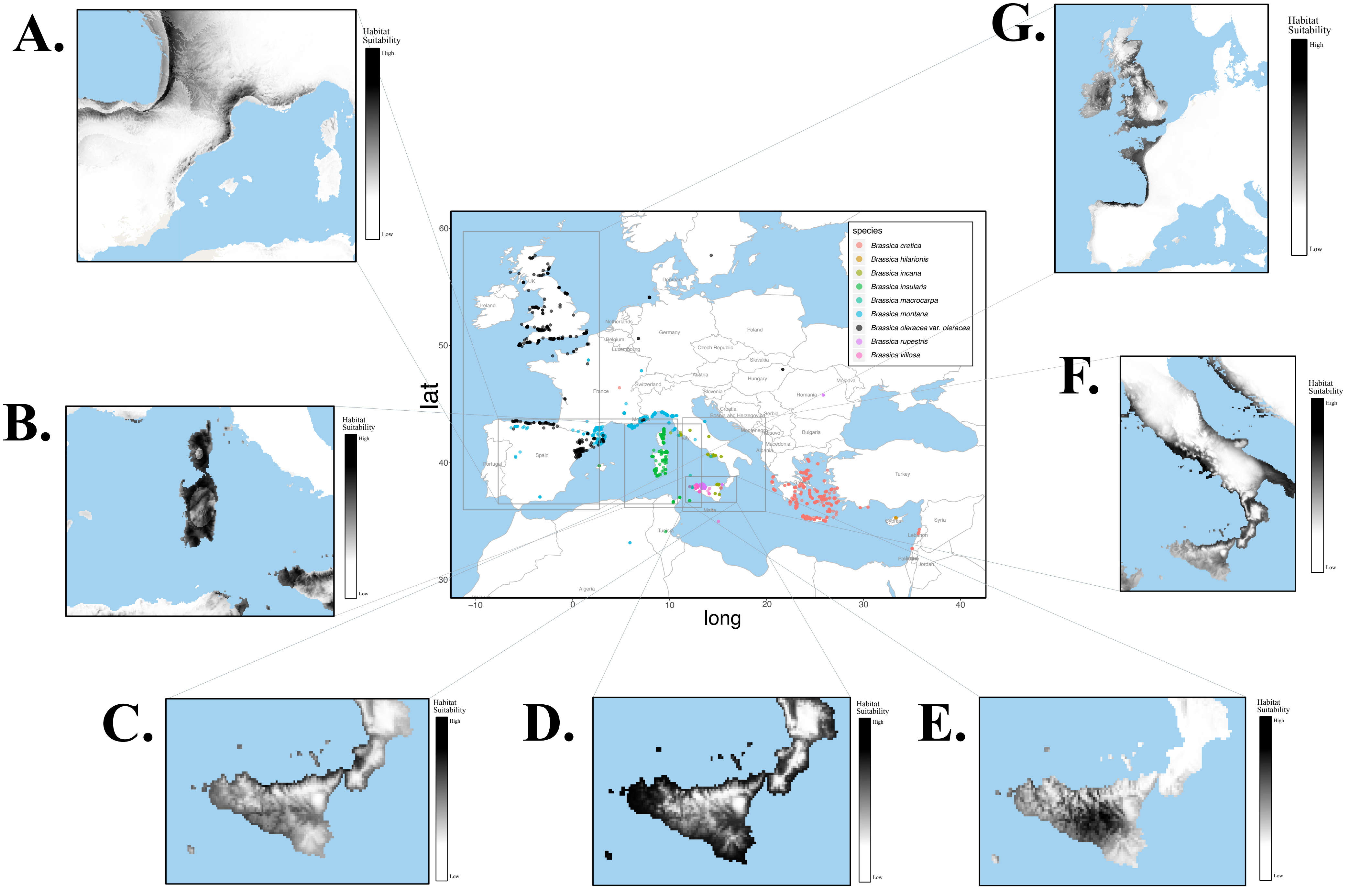
A.
B.
C.
D.
E.
F.
G.
Habitat Suitability
High
Low
species
Brassica cretica
Brassica hilarionis
Brassica incana
Brassica insularis
Brassica macrocarpa
Brassica montana
Brassica oleracea var. oleracea
Brassica rupestris
Brassica villosa
lat
long
60
50
40
30
−10
0
10
20
30
40
Ireland
UK
Denmark
Netherlands
Belgium
Luxembourg
Germany
Poland
Czech Republic
Slovakia
Austria
Hungary
Moldova
Romania
France
Switzerland
Slovenia
Croatia
Bosnia and Herzegovina
Serbia
Kosovo
Bulgaria
Macedonia
Albania
Spain
Portugal
Turkey
Malta
Syria
Lebanon
Jordan
Algeria

**A.** *B. oleracea* -175 Rep A, B, C, D

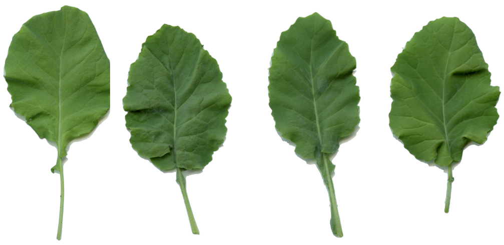

**B.** *B. oleracea* -176 Rep A, B, C, D

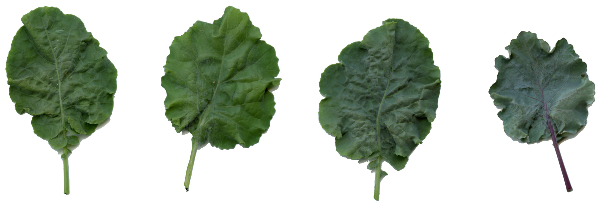

**C.** *B. oleracea* -177 Rep A, B, C, D

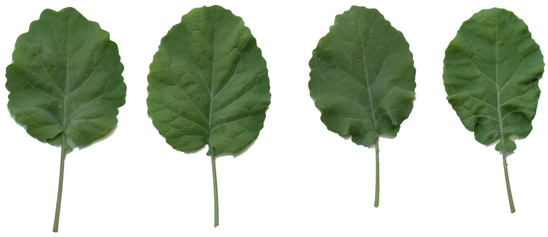

**D.** *B. cretica* -196 Rep A, B, C

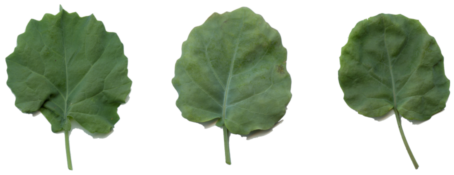

**E.** *B. cretica* -199 Rep A, B, C, D

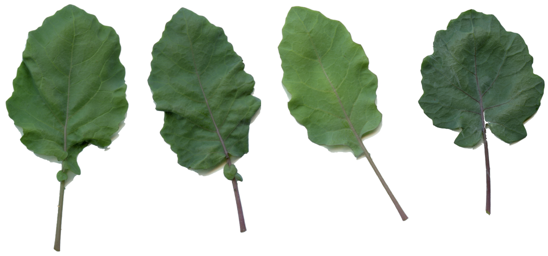

**F.** *B. montana* -222 Rep A, B, C, D

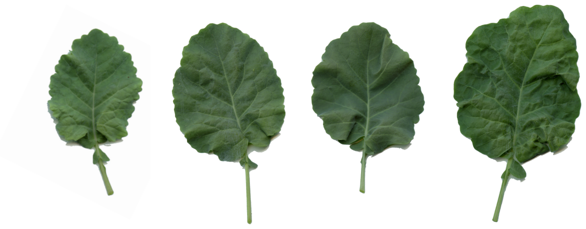

**G.** *B. incana* -204 Rep A, B, C, D

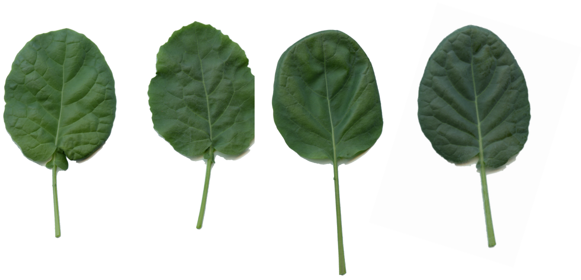

**H.** *B. incana* -207 Rep A, B, C, D

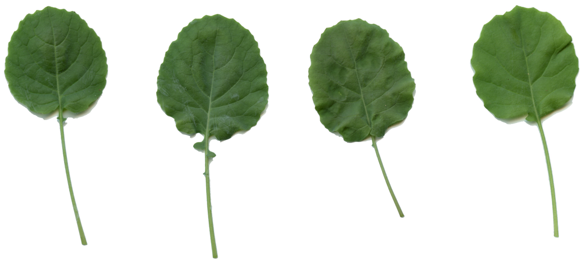

**A.** *B. cretica* -195 Rep A, B, C, D

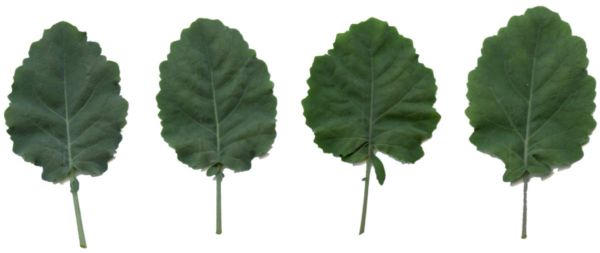

**B.** *B. incana* -205 Rep A, B, C, D

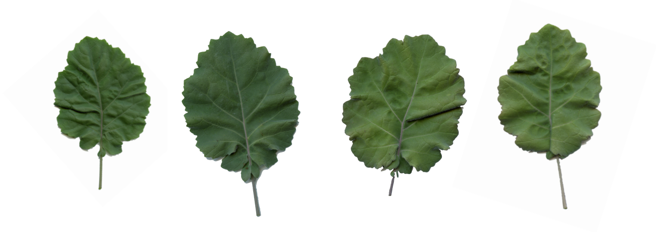

**C.** *B. incana* -208 Rep A, B, C, D

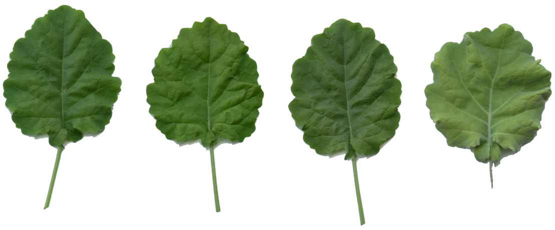

**D.** *B. incana* -209 Rep A, B, C, D

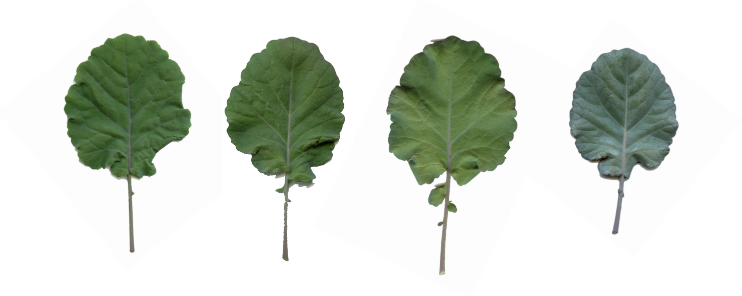

**E.** *B. villosa* -233 Rep A, B, C, D

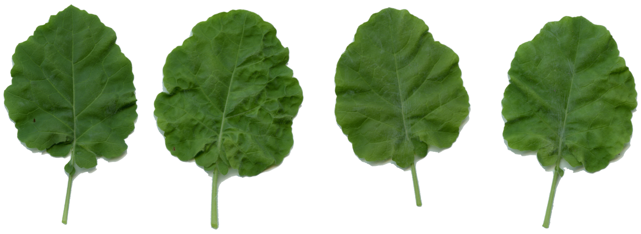

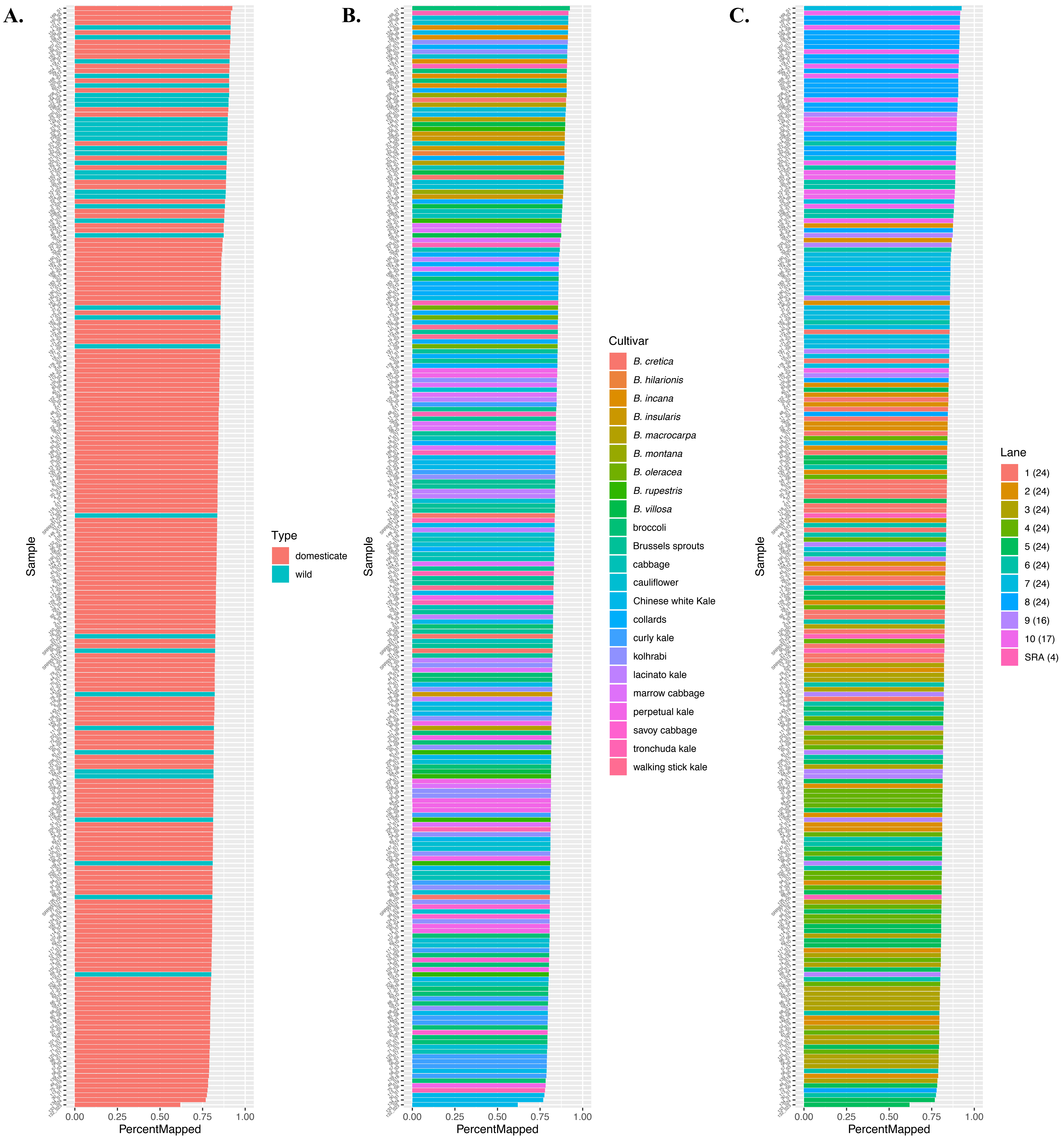

A.
B.
C.
Sample
PercentMapped
0.00
0.25
0.50
0.75
1.00
Type
domesticate
wild
Cultivar
B. cretica
B. hilarionis
B. incana
B. insularis
B. macrocarpa
B. montana
B. oleracea
B. rupestris
B. villosa
broccoli
Brussels sprouts
cabbage
cauliflower
Chinese white Kale
collards
curly kale
kolhrabi
lacinato kale
marrow cabbage
perpetual kale
savoy cabbage
tronchuda kale
walking stick kale
Lane
1 (24)
2 (24)
3 (24)
4 (24)
5 (24)
6 (24)
7 (24)
8 (24)
9 (16)
10 (17)
SRA (4)
